# Identification of type 2 diabetes loci in 433,540 East Asian individuals

**DOI:** 10.1101/685172

**Authors:** Cassandra N Spracklen, Momoko Horikoshi, Young Jin Kim, Kuang Lin, Fiona Bragg, Sanghoon Moon, Ken Suzuki, Claudia HT Tam, Yasuharu Tabara, Soo-Heon Kwak, Fumihiko Takeuchi, Jirong Long, Victor JY Lim, Jin-Fang Chai, Chien-Hsiun Chen, Masahiro Nakatochi, Jie Yao, Hyeok Sun Choi, Apoorva K Iyengar, Hannah J Perrin, Sarah M Brotman, Martijn van de Bunt, Anna L Gloyn, Jennifer E Below, Michael Boehnke, Donald W Bowden, John C Chambers, Anubha Mahajan, Mark I McCarthy, Maggie CY Ng, Lauren E Petty, Weihua Zhang, Andrew P Morris, Linda S Adair, Zheng Bian, Juliana CN Chan, Li-Ching Chang, Miao-Li Chee, Yii-Der Ida Chen, Yuan-Tsong Chen, Zhengming Chen, Lee-Ming Chuang, Shufa Du, Penny Gordon-Larsen, Myron Gross, Xiuqing Guo, Yu Guo, Sohee Han, Annie-Green Howard, Wei Huang, Yi-Jen Hung, Mi Yeong Hwang, Chii-Min Hwu, Sahoko Ichihara, Masato Isono, Hye-Mi Jang, Guozhi Jiang, Jost B Jonas, Yoichiro Kamatani, Tomohiro Katsuya, Takahisa Kawaguchi, Chiea-Chuen Khor, Katsuhiko Kohara, Myung-Shik Lee, Nannette R Lee, Liming Li, Jianjun Liu, Andrea O Luk, Jun Lv, Yukinori Okada, Mark A Pereira, Charumathi Sabanayagam, Jinxiu Shi, Dong Mun Shin, Wing Yee So, Atsushi Takahashi, Brian Tomlinson, Fuu-Jen Tsai, Rob M van Dam, Yong-Bing Xiang, Ken Yamamoto, Toshimasa Yamauchi, Kyungheon Yoon, Canqing Yu, Jian-Min Yuan, Liang Zhang, Wei Zheng, Michiya Igase, Yoon Shin Cho, Jerome I Rotter, Ya-Xing Wang, Wayne HH Sheu, Mitsuhiro Yokota, Jer-Yuarn Wu, Ching-Yu Cheng, Tien-Yin Wong, Xiao-Ou Shu, Norihiro Kato, Kyong-Soo Park, E-Shyong Tai, Fumihiko Matsuda, Woon-Puay Koh, Ronald CW Ma, Shiro Maeda, Iona Y Millwood, Juyoung Lee, Takashi Kadowaki, Robin G Walters, Bong-Jo Kim, Karen L Mohlke, Xueling Sim

**Affiliations:** Department of Genetics, University of North Carolina at Chapel Hill, Chapel Hill, NC, USA; Laboratory for Endocrinology, Metabolism and Kidney Diseases, RIKEN Centre for Integrative Medical Sciences, Yokohama, Japan; Division of Genome Research, Center for Genome Science, National Institute of Health, Cheongjusi, Korea; Nuffield Department of Population Health, University of Oxford, Oxford, UK; Laboratory for Statistical and Translational Genetics, RIKEN Centre for Integrative Medical Sciences, Yokohama, Japan; Department of Diabetes and Metabolic Diseases, Graduate School of Medicine, The University of Tokyo, Tokyo, Japan; Department of Statistical Genetics, Osaka University Graduate School of Medicine, Osaka, Japan; Department of Medicine and Therapeutics, The Chinese University of Hong Kong, Hong Kong, China; Chinese University of Hong Kong-Shanghai Jiao Tong University Joint Research Centre in Diabetes Genomics and Precision Medicine, The Chinese University of Hong Kong, Hong Kong, China; Center for Genomic Medicine, Kyoto University Graduate School of Medicine, Kyoto, Japan; Department of Internal Medicine, Seoul National University Hospital, Seoul, South Korea; Department of Gene Diagnostics and Therapeutics, Research Institute, National Center for Global Health and Medicine, Tokyo, Japan; Division of Epidemiology, Department of Medicine, Vanderbilt University Medical Center, Nashville, TN, USA; Saw Swee Hock School of Public Health, National University of Singapore and National University Health System, Singapore, Singapore; Institute of Biomedical Sciences, Academia Sinica, Taipei, Taiwan; Department of Nursing, Nagoya University Graduate School of Medicine, Nagoya University Hospital, Nagoya, Japan; Department of Pediatrics, The Institute for Translational Genomics and Population Sciences, LABioMed at Harbor-UCLA Medical Center, Torrance, CA, USA; Biomedical Science, Hallym University, Chuncheon, South Korea; Oxford Centre for Diabetes, Endocrinology and Metabolism, University of Oxford, Oxford, UK; Wellcome Centre for Human Genetics, University of Oxford, Oxford, UK; Oxford NIHR Biomedical Research Centre, Oxford University Hospitals NHS Foundation Trust, Churchill Hospital, Oxford, UK; Vanderbilt Genetics Institute, Division of Genetic Medicine, Vanderbilt University Medical Center, Nashville, TN, USA; Human Genetics Center, School of Public Health, The University of Texas Health Science Center at Houston, Houston, TX, USA; Department of Biostatistics and Center for Statistical Genetics, University of Michigan, Ann Arbor, MI, USA; Center for Genomics and Personalized Medicine Research, Center for Diabetes Research, Wake Forest School of Medicine, Winston-Salem, NC, USA; Department of Biochemistry, Wake Forest School of Medicine, Winston-Salem, NC, USA; Lee Kong Chian School of Medicine, Nanyang Technological University, Singapore, Singapore; Department of Epidemiology and Biostatistics, Imperial College London, London, UK; Department of Cardiology, Ealing Hospital, London North West Healthcare NHS Trust, Middlesex, UK; Imperial College Healthcare NHS Trust, Imperial College London, London, UK; MRC-PHE Centre for Environment and Health, Imperial College London, London, UK; Department of Biostatistics, University of Liverpool, Liverpool, UK; School of Biological Sciences, University of Manchester, Manchester, UK; Department of Nutrition, Gillings School of Global Public Health, University of North Carolina at Chapel Hill, Chapel Hill, NC, USA; Chinese Academy of Medical Sciences, Beijing, China; Hong Kong Institute of Diabetes and Obesity, The Chinese University of Hong Kong, Hong Kong, China; Li Ka Shing Institute of Health Sciences, The Chinese University of Hong Kong, Hong Kong, China; Singapore Eye Research Institute, Singapore National Eye Centre, Singapore, Singapore; Division of Endocrinology & Metabolism, Department of Internal Medicine, National Taiwan University Hospital, Taipei, Taiwan; Institute of Preventive Medicine, School of Public Health, National Taiwan University, Taipei, Taiwan; Department of Laboratory Medicine and Pathology, University of Minnesota, Minneapolis, MN, USA; Department of Biostatistics, Carolina Population Center, Gillings School of Global Public Health, University of North Carolina at Chapel Hill, Chapel Hill, NC, USA; Department of Genetics, Shanghai-MOST Key Laboratory of Health and Disease Genomics, Chinese National Human Genome Center at Shanghai, Shanghai, China; Division of Endocrine and Metabolism, Tri-Service General Hospital Songshan Branch, Taipei, Taiwan; School of Medicine, National Defense Medical Center, Taipei, Taiwan; Section of Endocrinology and Metabolism, Department of Medicine, Taipei Veterans General Hospital, Taipei, Taiwan; School of Medicine, National Yang-Ming University, Taipei, Taiwan; Department of Environmental and Preventive Medicine, Jichi Medical University School of Medicine, Shimotsuke, Japan; Department of Ophthalmology, Medical Faculty Mannheim of the University of Heidelberg, Mannheim, Germany; Laboratory of Complex Trait Genomics, Department of Computational Biology and Medical Sciences, Graduate School of Frontier Sciences, The University of Tokyo, Tokyo, Japan; Department of Clinical Gene Therapy, Osaka University Graduate School of Medicine, Osaka, Japan; Department of Geriatric and General Medicine, Graduate School of Medicine, Osaka University, Osaka, Japan; Genome Institute of Singapore, Agency for Science, Technology and Research, Singapore, Singapore; Department of Biochemistry, National University of Singapore, Singapore, Singapore; Department of Regional Resource Management, Ehime University Faculty of Collaborative Regional Innovation, Ehime, Japan; Severance Biomedical Science Institute and Department of Internal Medicine, Yonsei University College of Medicine, Seoul, South Korea; Department of Anthropology, Sociology and History, University of San Carlos, Cebu City, Philippines; Departments of Epidemiology & Biostatistics, Peking University Health Science Centre, Peking University, Beijing, China; Department of Medicine, Yong Loo Lin School of Medicine, National University of Singapore and National University Health System, Singapore, Singapore; Laboratory of Statistical Immunology, Immunology Frontier Research Center (WPI-IFReC), Osaka University, Osaka, Japan; Division of Epidemiology and Community Health, School of Public Health, University of Minnesota, Minneapolis, MN, USA; Ophthalmology & Visual Sciences Academic Clinical Program (Eye ACP), Duke-NUS Medical School, Singapore, Singapore; Department of Ophthalmology, Yong Loo Lin School of Medicine, National University of Singapore and National University Health System, Singapore, Singapore; Department of Genomic Medicine, National Cerebral and Cardiovascular Center, Osaka, Japan; Department of Medical Genetics and Medical Research, China Medical University Hospital, Taichung, Taiwan; Department of Nutrition, Harvard T.H Chan School of Public Health, Boston, MA, USA; State Key Laboratory of Oncogene and Related Genes & Department of Epidemiology, Shanghai Cancer Institute, Renji Hospital, Shanghai Jiaotong University School of Medicine, Shanghai, China; Department of Medical Biochemistry, Kurume University School of Medicine, Kurume, Japan; Division of Cancer Control and Population Sciences, UPMC Hillman Cancer Center, University of Pittsburgh, Pittsburgh, PA, USA; Department of Epidemiology, Graduate School of Public Health, University of Pittsburgh, Pittsburgh, PA, USA; Department of Anti-aging Medicine, Ehime University Graduate School of Medicine, Ehime, Japan; Departments of Pediatrics and Medicine, The Institute for Translational Genomics and Population Sciences, LA BioMed at Harbor-UCLA Medical Center, Torrance, CA, USA; Beijing Institute of Ophthalmology, Ophthalmology and Visual Sciences Key Laboratory, Beijing Tongren Hospital, Capital Medical University, Beijing, China; Division of Endocrinology and Metabolism, Department of Medicine, Taichung Veterans General Hospital, Taichung, Taiwan; Kurume University School of Medicine, Kurume, Japan; Department of Internal Medicine, Seoul National University College of Medicine, Seoul, South Korea; Department of Molecular Medicine and Biopharmaceutical Sciences, Graduate School of Convergence Science and Technology, Seoul National University, Seoul, South Korea; Duke-NUS Medical School, Singapore, Singapore; Health Services and Systems Research, Duke-NUS Medical School, Singapore, Singapore; Department of Advanced Genomic and Laboratory Medicine, Graduate School of Medicine, University of the Ryukyus, Okinawa, Japan; Division of Clinical Laboratory and Blood Transfusion, University of the Ryukyus Hospital, Okinawa, Japan; Medical Research Council Population Health Research Unit, University of Oxford, Oxford, UK

## Abstract

Meta-analyses of genome-wide association studies (GWAS) have identified >240 loci associated with type 2 diabetes (T2D), however most loci have been identified in analyses of European-ancestry individuals. To examine T2D risk in East Asian individuals, we meta-analyzed GWAS data in 77,418 cases and 356,122 controls. In the main analysis, we identified 298 distinct association signals at 178 loci, and across T2D association models with and without consideration of body mass index and sex, we identified 56 loci newly implicated in T2D predisposition. Common variants associated with T2D in both East Asian and European populations exhibited strongly correlated effect sizes. New associations include signals in/near *GDAP1*, *PTF1A*, *SIX3, ALDH2,* a microRNA cluster, and genes that affect muscle and adipose differentiation. At another locus, eQTLs at two overlapping T2D signals act through two genes, *NKX6-3* and *ANK1*, in different tissues. Association studies in diverse populations identify additional loci and elucidate disease genes, biology, and pathways.

Type 2 diabetes (T2D) is a common metabolic disease primarily caused by insufficient insulin production and/or secretion by the pancreatic β cells and insulin resistance in peripheral tissues^1^. Most genetic loci associated with T2D have been identified in populations of European (EUR) ancestry, including a recent meta-analysis of genome-wide association studies (GWAS) of nearly 900,000 individuals of European ancestry that identified >240 loci influencing the risk of T2D^2^. Differences in allele frequency between ancestries affect the power to detect associations within a population, particularly among variants rare or monomorphic in one population but more frequent in another^3,4^. Although smaller than studies in European populations, a recent T2D meta-analysis in almost 200,000 Japanese individuals identified 28 additional loci^4^. The relative contributions of different pathways to the pathophysiology of T2D may also differ between ancestry groups. For example, in East Asian (EAS) populations, T2D prevalence is greater than in European populations among people of similar body mass index (BMI) or waist circumference^5^. We performed the largest meta-analysis of East Asian individuals to identify new genetic associations and provide insight into T2D pathogenesis.

## RESULTS

We conducted a fixed-effect inverse-variance weighted GWAS meta-analysis combining 23 studies imputed to the 1000 Genomes Phase 3 reference panel from the Asian Genetic Epidemiology Network (AGEN) consortium (Supplementary Tables 1-3). We performed sex-combined T2D association without BMI adjustment in 77,418 T2D cases and 356,122 controls (effective sample size, N_eff_=211,793) and with BMI adjustment in 54,481 T2D cases and 224,231 controls (N_eff_= 135,780). In the set of studies with BMI-adjusted analyses, we also tested for T2D association in models stratified by sex (Supplementary Figure 1). We defined “lead” variants as the strongest T2D-associated variants with *P*<5×10^−8^ and defined the region +/− 500 kb from the lead variant as a locus. A locus was considered novel if the lead variant was located at least 500 kb away from previously reported T2D-associated variants in any ancestry.

Using summary association statistics for ~11.7 million variants without adjustment for BMI (Supplementary Figure 1; Supplementary Tables 1-3), we identified lead variants at 178 loci to be associated with T2D, of which 49 were novel (Table 1; Supplementary Figure 2; Supplementary Table 4). Lead variants at all novel loci were common (MAF≥5%; Supplementary Figure 3), except for two low-frequency lead variants near *GDAP1* (MAF=2.4%), which regulates mitochondrial proteins and metabolic flux in skeletal muscle^6^, and *PTF1A* (MAF=4.7%), which encodes a transcription factor required for pancreatic acinar cell development^7^. Lead variants met a stricter *P*-value threshold for significance based on Bonferroni correction for 11.7 million tests (*P*<4.3×10^−9^) at 147 of the 178 loci, including 31 of the 49 novel loci.

**Table 1.**
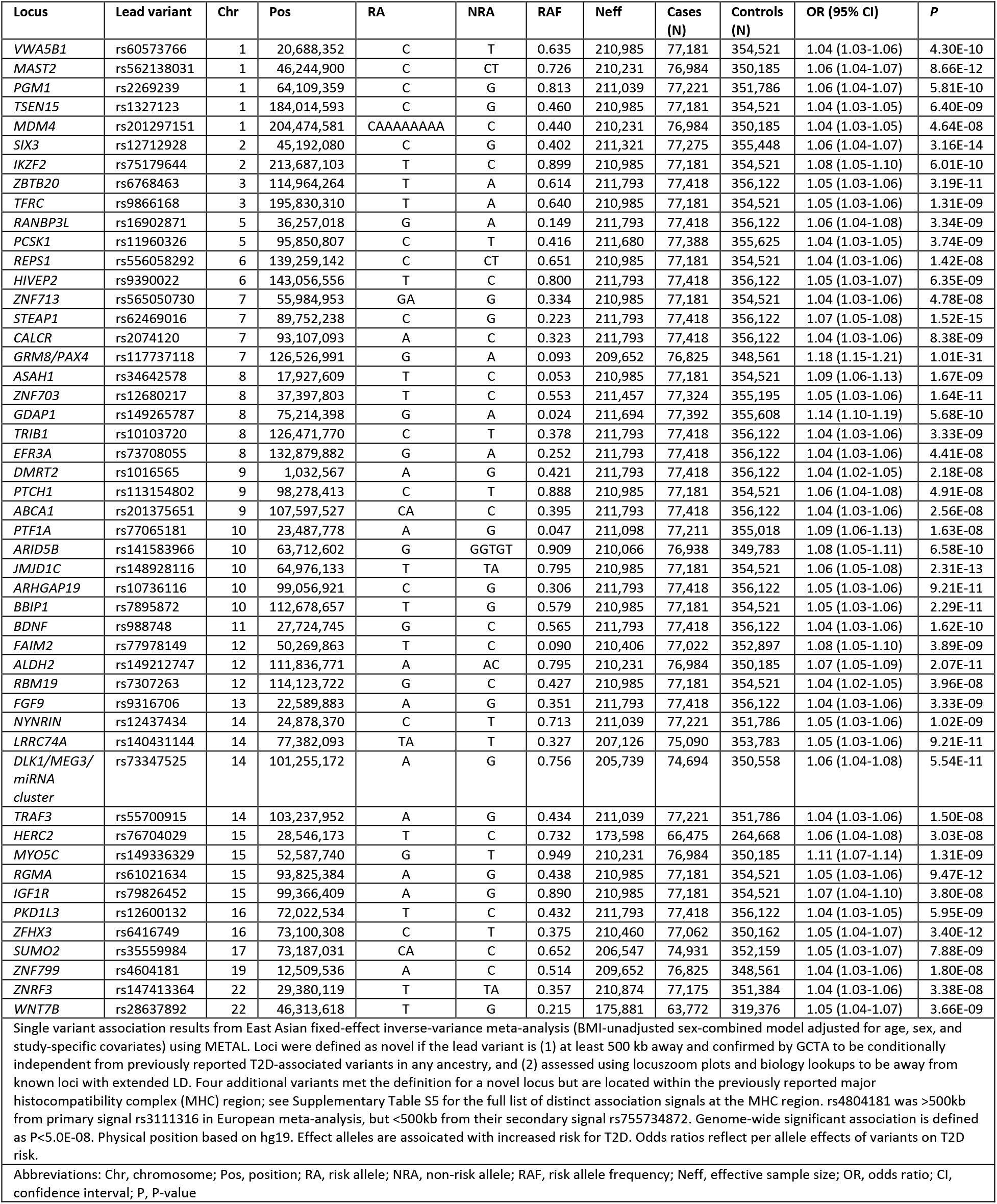
Novel lead variants associated with type 2 diabetes in East Asians.

Using GCTA^8^, we identified 298 distinct signals that met a locus-wide significance threshold of *P*<1×10^−5^ (Supplementary Table 5), 204 of which were genome-wide significant (*P*<5×10^−8^). Overall, we observed 2-4 signals at 50 loci and ≥5 signals at 11 loci. Among the 49 novel loci, 9 loci had two signals and the locus at *WNT7B* had three signals. Among the ten loci with the most significant meta-analysis *P-*values of association, seven contained ≥5 distinct signals (16 signals at *INS*/*IGF2*/*KCNQ1*; 7 signals at *CDKN2A/B* and *GRM8*/*PAX4*/*LEP*; 5 signals at *CDKAL1*, *HHEX*/*IDE, CDC123*/*CAMK1D,* and *TCF7L2;* Supplementary Figure 4; Supplementary Tables 5). The seven distinct association signals at the *GRM8*/*PAX4*/*LEP* locus span 1.4 Mb, and no evidence of T2D association at this locus has yet been reported in non-East Asian ancestry groups^2,9^ (Supplementary Figure 4C). Joint analyses confirmed independent associations (LD *r*^2^=0.0025) at two previously reported *PAX4* missense variants^10^, rs2233580 [Arg192His: risk allele frequency (RAF)=8.6%, OR=1.31, 95% CI 1.27 – 1.34, *P*_GCTA_=3.0×10^−89^] and rs3824004 (Arg192Ser: RAF=3.4%, OR=1.23, 95% CI 1.19-1.28, *P*_GCTA_=4.3×10^−29^). The association signals at this locus also include variants near *LEP*, which encodes leptin, a hormone that regulates appetite^11^; increased leptin levels are associated with obesity and T2D, with greater increase in leptin levels per unit of BMI in Chinese individuals compared to those of African-American, European, and Hispanic ancestries^12^.

At the previously reported *ANK1/NKX6-3* locus^2,13,14^, we observed three distinct T2D association signals, two of which overlap and consist of variants spanning only ~25 kb (Figure 1). Given conflicting interpretation of candidate genes^2,15,16^, we compared the T2D-association signals identified in East Asian individuals to eQTLs reported at this locus in islets^2,16–18^, subcutaneous adipose^19^, and skeletal muscle^15^. At the strongest signal, the lead T2D-associated variant, rs33981001, is in high LD with the lead *cis*-eQTL variant for *NKX6-3* in pancreatic islets (rs12549902; EAS LD *r*^2^=0.79, EUR *r*^2^=0.83)^16^, and the T2D risk allele is associated with decreased expression of *NKX6-3* (β=-0.36, *P*=6.1×10^−7^; Figure 1)^20^. *NKX6-3*, or NK6 homeobox 3, encodes a pancreatic islet transcription factor required for the development of alpha and β cells in the pancreas^21^ and has been shown to influence insulin secretion^16^. At the second T2D-association signal, rs62508166 is in high LD with the lead *cis*-eQTL variant for *ANK1* in subcutaneous adipose tissue^19^ and skeletal muscle^15^ (rs516946; EAS LD *r*^2^=0.96, EUR *r*^2^=0.80), and the T2D risk allele is associated with increased expression of *ANK1* (subcutaneous adipose: β=0.20, *P*=1.8×10^−7^; skeletal muscle: β=1.01, *P*=2.8×10^−22^). *ANK1* belongs to the ankyrin family of integral membrane proteins that has been shown to affect glucose uptake in skeletal muscle, and changes in expression levels may lead to insulin resistance^22^. Together, these GWAS and eQTL signals suggest that variants within this ~25 kb region act to increase or decrease expression levels of two different genes in different tissues to increase T2D risk.

**Figure 1:**
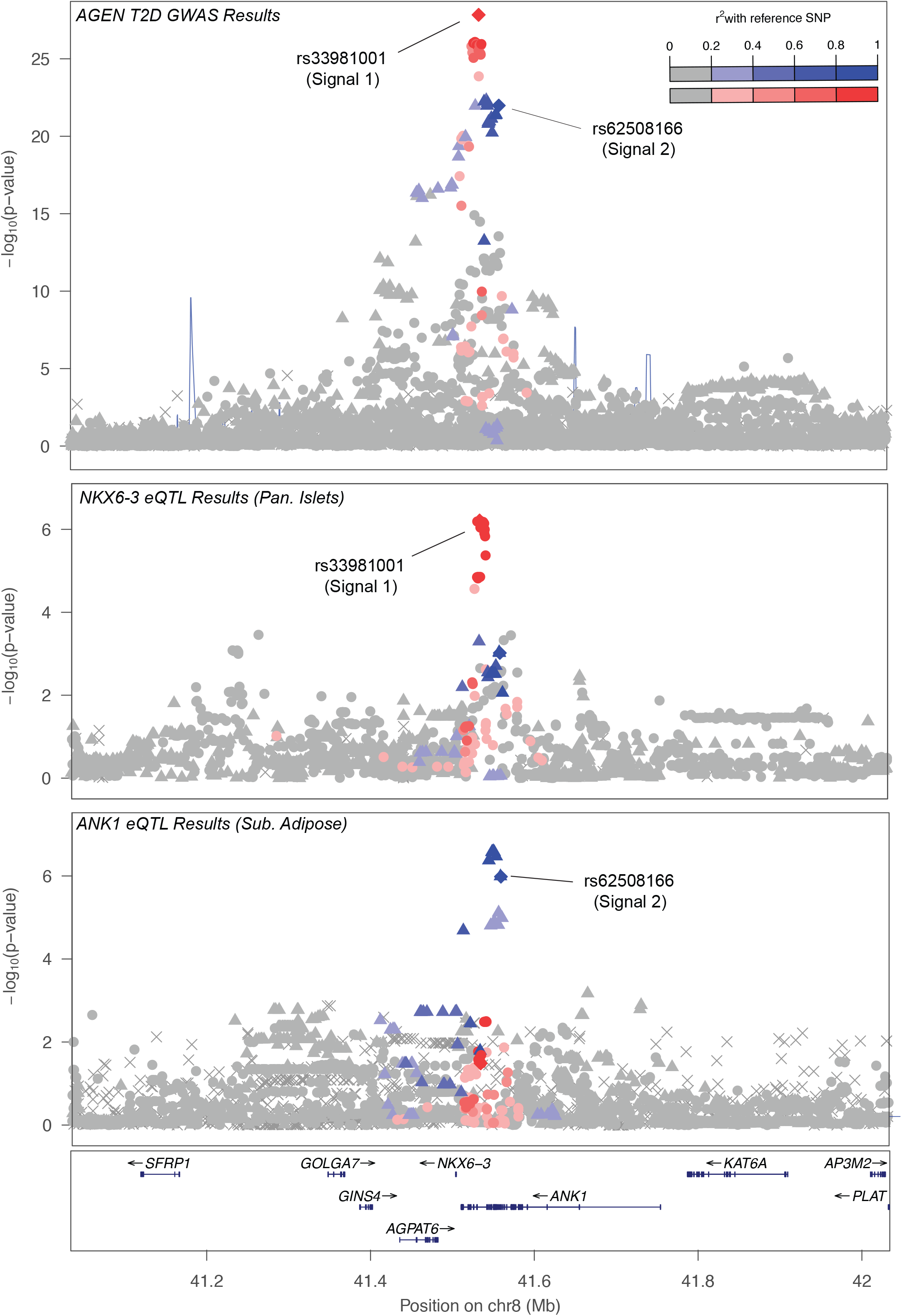
Two distinct T2D-association signals at the *ANK1-NKX6-3* locus associated with expression levels of two transcripts in two tissues. (A) Regional association plot for East Asian sex-combined BMI-unadjusted meta-analysis at *ANK1-NKX6-3* locus. Approximate conditional analysis using GTCA identified three distinct T2D-association signals at this locus (signal 1, rs33981001; signal 2, rs62508166; signal 3, rs144239281, in order of strength of association). Using 1000G Phase3 East Asian LD, variants are colored in red and blue with the first and second distinct signals respectively (lead variants represented as diamonds). (B) Variant rs12549902, in high LD (EAS LD *r*^2^=0.80, EUR *r*^2^=0.83) with T2D signal 1, shows the strongest association with expression levels of *NXK6-3* in pancreatic islets in 118 individuals^16^. (C) Variant rs516946, in high LD (EAS LD *r*^2^=0.96, EUR *r*^2^=0.80) with T2D signal 2, shows the strongest association with expression levels of *ANK1* in subcutaneous adipose tissue in 770 individuals^19^. As rs62508166 is not available in the subcutaneous adipose tissue data set, a variant in perfect LD (rs28591316) was used and is represented by the blue diamond variant.

In T2D association analyses adjusted for BMI, we identified an additional ten loci, four of which were not reported previously for T2D, including loci near *MYOM3/SRSF10*, *TSN*, *GRB10*, and *NID2* (Supplementary Figure 5A; Supplementary Table 4). At the *NID2* locus, the T2D risk allele is associated with lower BMI in East Asian individuals, consistent with a lipodystrophy phenotype^23,24^. Among the combined 188 loci identified in models with and without adjustment for BMI, effect sizes were highly correlated (Pearson correlation r=0.99; Supplementary Figures 1 and 6). The locus with the strongest heterogeneity between the two models was *FTO* (*P*_het_=5.6×10^−3^), although even this locus failed to surpass a Bonferroni-corrected threshold for significant heterogeneity (*P*_het_<0.05/188=2.7×10^−4^).

In sex-stratified analyses of males (28,027 cases and 89,312 controls) and females (27,370 cases and 135,055 controls), we identified one additional novel male-specific locus near *IFT81* and two additional novel female-specific loci near *CPS1* and *LMTK2* (Supplementary Figure 5B and 5C; Supplementary Table 6). The lead *CPS1* variant rs1047891 (Thr1412Asn) has been reported previously to have a stronger effect in females than in males for cardiovascular disease and several blood metabolites^25^. Taken together, we identified a total of 56 novel loci across BMI-unadjusted, BMI-adjusted, and sex-stratified models.

Among all T2D-associated loci, a region spanning almost 2 Mb on chromosome 12 near *ALDH2* exhibited the strongest differences between sexes (rs12231737, *P*_het_=2.6×10^−19^), with compelling evidence of association in males (*P*=5.5×10^−27^) and no evidence for association in females (*P*=0.19) (Supplementary Figure 7; Supplementary Table 6). Further, joint conditional analyses revealed two conditionally distinct signals (rs12231737, *P*_GCTA_=1.7×10^−21^; rs557597782, *P*_GCTA_=4.7×10^−7^) in males only. *ALDH2* encodes aldehyde dehydrogenase 2 family member, a key enzyme in alcohol metabolism that converts acetaldehyde into acetic acid. This stretch of T2D associations in males reflects a long LD block that arose due to a recent selective sweep in East Asian individuals and results in flushing, nausea, and headache following alcohol consumption^26^. The most significantly associated missense variant in moderate LD with rs12231737 (*r*^2^=0.68) was common functional variant rs671 (*ALDH2* Glu504Lys: RAF=77.4%, OR=1.16, 95% CI 1.15 – 1.18, *P*_males_=4.2×10^−24^), which leads to reduced ALDH2 activity and reduced alcohol metabolism, and have previously been reported to be associated with cardiometabolic traits in East Asian populations; the T2D risk allele is associated with better tolerance for alcohol and increased BMI, increased systolic and diastolic blood pressure, and increased triglycerides, but increased high-density lipoprotein, decreased low-density lipoprotein, and decreased cardiovascular risk^27-31^. The strong sexual dimorphism observed at this locus may be due to differences in alcohol consumption patterns between males and females^27,29^ and/or differences in the effect of alcohol on insulin sensitivity^32^.

With an effective sample size comparable to the largest study of T2D in European individuals (East Asian N_eff_=211,793; European N_eff_ = 231,436)^2^ and imputation to a dense 1000 Genomes reference panel, our results provide the most comprehensive and precise catalogue of East Asian T2D effects to date for comparisons across ancestries (Figure 2; Supplementary Table 7). For 178 EAS T2D loci and 231 EUR T2D loci (unadjusted for BMI) identified in a European meta-analysis^2^, we compared the per-allele effect sizes for the 343 variants available in both datasets (i.e. polymorphic and passed quality control), including lead variants from both ancestries at shared signals. Overall, the per-allele effect sizes between the two ancestries were moderately correlated (r=0.54; Figure 2A). When the comparison was restricted to the 290 variants that are common (MAF≥5%) in both ancestries, the effect size correlation increased to r=0.71 (Figure 2B; Supplementary Figure 8). This effect size correlation further increased to r=0.88 for 116 variants significantly associated with T2D (*P*<5×10^−8^) in both ancestries. While the overall effect sizes for all 343 variants appear, on average, to be stronger in East Asian individuals than European, this trend is reduced when each locus is represented only by the lead variant from one population (Supplementary Figure 9). Specifically, many variants identified with larger effect sizes in the European meta-analysis are missing from the comparison because they were rare/monomorphic or poorly imputed in the East Asian meta-analysis, for which imputation reference panels are less comprehensive compared to the European-centric Haplotype Reference Consortium panel.

**Figure 2:**
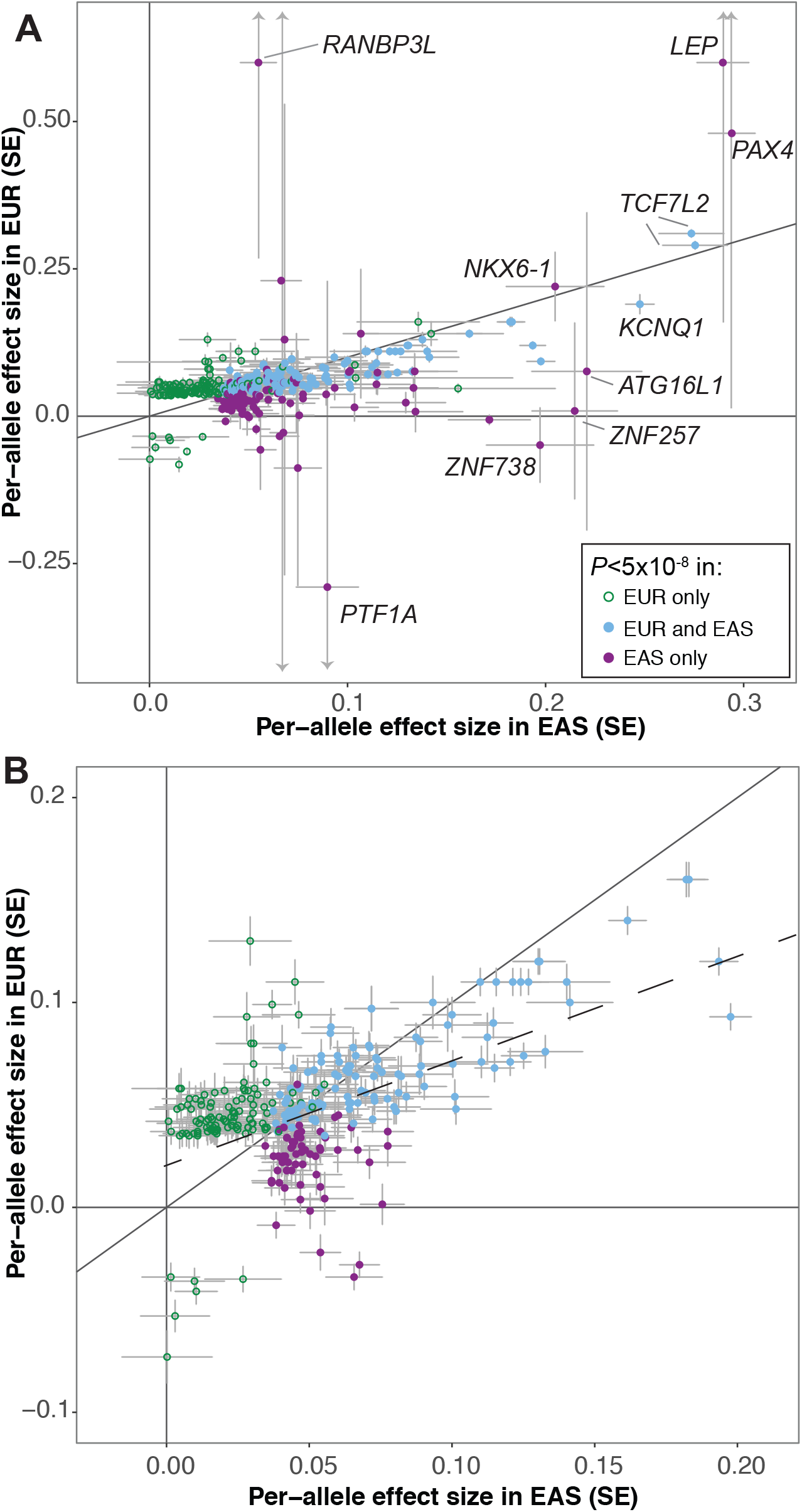
Effect size comparison of lead variants identified in this East Asian T2D GWAS BMI-unadjusted meta-analysis and previous European T2D GWAS meta-analysis^2^. For 343 lead variants identified from the two BMI unadjusted meta-analyses, per-allele effect sizes (β) from the European meta-analysis (*y*-axis) were plotted against per-allele effect sizes from this East Asian meta-analysis (*x*-axis). (A) All 343 lead variants; (B) 290 lead variants with minor allele frequency ≥5% in both ancestries. Variants are colored purple if they were significant (*P*<5×10^−8^) in the East Asian analysis only, green if they were significant in European analysis only, and blue if they were significant in both the East Asian and European analysis (see Methods and Supplementary Table 7). The dashed diagonal line represents the trend line across all plotted variants.

Variants exhibiting the largest differences in effect sizes across ancestries are generally rare (MAF ≤0.1%) in European populations but common (e.g. *PAX4*, *RANBP3L*) or low frequency (e.g. *ZNF257*, *DGKD*) in East Asian populations. For example, rs142395395 near *ZNF257* (RAF=96.9%, OR=1.24, 95% CI 1.19-1.29, *P*=7.0×10^−23^) has been reported only twice in 15,414 individuals of non-Finnish European ancestry from the gnomAD database^33^. This variant tags a previously described inversion of 415 kb observed only in East Asian individuals that disrupts the coding sequence and expression of *ZNF257,* as well as lymphoblastoid expression of 81 downstream genes and transcripts^34^. These data suggest that *ZNF257* and/or downstream target genes influence T2D susceptibility (Supplementary Figure 10).

We identified many loci for which the lead variants exhibited similar allele frequencies and effect sizes in both the East Asian and European meta-analyses, but only reached genome-wide significance in the East Asian meta-analysis. Given shared susceptibility across ancestry groups, these loci may be detected in non-East Asian populations when sample sizes increase. Among these variants is rs117624659, located near *NKX6-1* (*P*_EAS_ = 2.0×10^−16^, *P*_EUR_ =2.2×10^−4^). This lead variant overlaps a highly conserved region that shows open chromatin specific to pancreatic islets. We conducted transcriptional reporter assays in MIN6 mouse insulinoma cells and observed that rs117624659 exhibited significant allelic differences in enhancer activity (Figure 3; Supplementary Figure 11). In the pancreas, NK6 homeobox 1 (NKX6.1) is required for the development of insulin-producing β cells and is a potent bifunctional transcriptional regulator^35^. Further, inactivation of *Nkx6.1* in mice demonstrated rapid-onset diabetes due to defects in insulin biosynthesis and secretion^36^. Unexpectedly, the T2D risk allele showed increased transcriptional activity, suggesting that the variant does not act in isolation or that *NXK6-1* is not the target gene.

**Figure 3:**
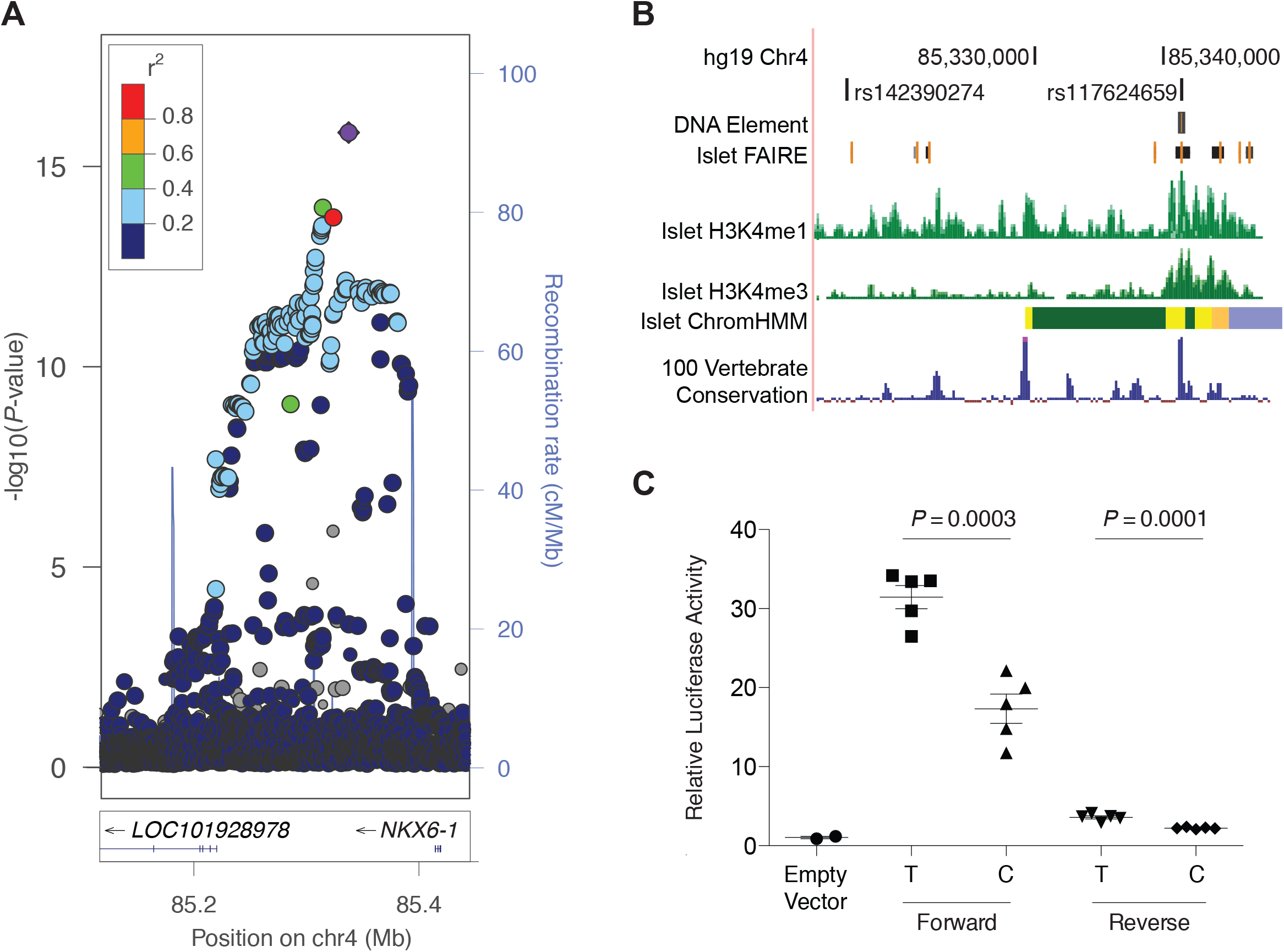
rs117624659 at *NKX6-1* locus exhibits allelic differences in transcriptional activity. (A) rs117624659 (purple diamond) shows the strongest association with T2D in the region. Variants are colored based on 1000G Phase 3 East Asian LD with rs117624659. (B) rs117624659 and an additional candidate variant rs142390274 in high pairwise LD (*r*^2^>0.80) span a 22 kb region approximately 75 kb upstream of *NKX6-1*. rs117624659 overlaps a region of open chromatin in pancreatic islets and lies within a region conserved across vertebrates. (C) rs117624659-T, associated with increased risk of T2D, showed greater transcriptional activity in an element cloned in both forward and reverse orientations with respect to *NKX6-1* in MIN6 cells compared to rs117624659-C and an “empty vector” containing a minimal promoter.

At one of the novel T2D-associated loci near *SIX3*, the risk allele of East Asian lead variant rs12712928 (RAF=40.2%, OR=1.06, 95% CI 1.04 – 1.07, *P*=3.2×10^−14^) is common across non-East Asian ancestries, ranging from 16.0% in Europeans to 26.4% in South Asians; however, there was no evidence of association in the other ancestry groups (meta-analysis: OR=0.98, 95% CI 0.96 – 0.99, *P*=2.9×10^−3^) (Figure 4A; Supplementary Figure 12; Supplementary Table 8). Within the East Asian meta-analysis, the direction of effect is consistent across East Asian countries (Figure 4B) and within the contributing cohorts (Supplementary Figure 13). The T2D risk allele rs12712928-C is associated with higher fasting glucose levels in East Asian populations^37,38^, has the strongest association with lower expression levels of both *SIX3* and *SIX2* in pancreatic islets^17^, and demonstrated allele-specific binding to the transcription factor GABPA and significantly lower levels of transcriptional activity^38^. While larger studies in other ancestry groups could improve the accuracy of the effect estimate, current evidence suggests that the T2D association near *SIX3* is specific to East Asian populations.

**Figure 4:**
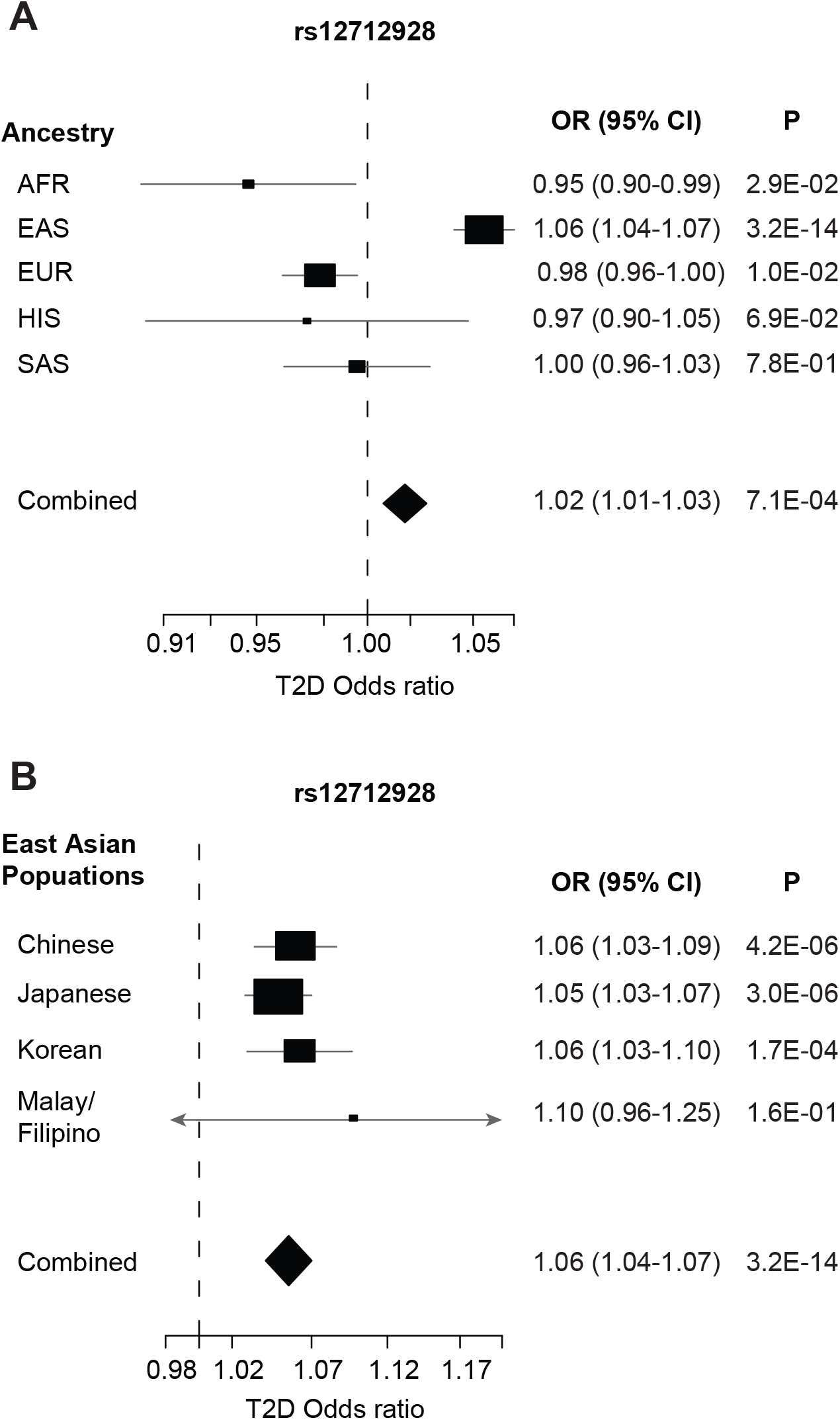
Forest plots of BMI-unadjusted meta-analysis association results at *SIX3*-*SIX2* locus. Odds ratios (black boxes) and 95% confidence intervals (horizontal lines) for T2D associations at the lead East Asian variant (rs12721928) are presented (A) across ancestries of African-American (AFR), East Asian (EAS), European (EUR)^2^, Hispanic (HIS), and South Asian (SAS) individuals, and (B) within four major East Asian populations (Chinese, Japanese, Korean, and Malay/Filipino combined due to small sample sizes). The size of the box is proportional to the sample size of each contributing population.

To identify potential candidate genes underlying the T2D-association signals identified in East Asian individuals, we further characterized 88 loci, including known and novel loci, for which the lead variant at the primary East Asian association signal is located >500 kb from the lead variant of any primary European T2D association signal^2^ (Supplementary Table 9). We characterized loci using prior trait associations, *cis*-regulatory effects on expression (colocalized eQTL), predicted effects on protein sequence, and a literature search (Supplementary Tables 10-13). Based on association results from cardiometabolic trait consortia^39^, Biobank Japan^40^, and the UK Biobank^41^, the lead T2D-associated variant at 19 of the 88 loci was associated (*P*<5×10^−8^) with at least one additional cardiometabolic trait, most frequently BMI or a fat mass-related trait (16 loci; Supplementary Tables 10 and 12). At 12 of the examined loci, T2D signals were colocalized with *cis*-eQTLs for transcripts in subcutaneous adipose tissue (n=5), skeletal muscle (n=3), pancreas (n=2), pancreatic islets (n=3), or whole blood (n=5; Supplementary Tables 11-12). At 19 loci, the lead T2D-associated variant or a variant in high LD with it (East Asian *r*^2^>0.80) alter the protein sequence (Supplementary Tables 12). These variants affect mesenchymal stem cell differentiation and adipogenesis (*GIT2*, *STEAP2* and *JMJD1C*), muscle stem cell biology (*CALCR*), glucose metabolism (*PGM1* and *SCTR*), and insulin secretion (*FGFR4*; Supplementary Table 13). While mechanistic inference is required, these potential molecular mechanisms suggest new T2D susceptibility genes primarily detected by analyses in East Asians.

T2D loci were also identified at clusters of noncoding RNAs with roles in islet β cell function. One locus includes a set of microRNAs specifically expressed in islet β cells, the maternally expressed noncoding RNA *MEG3,* and the paternally expressed gene *DLK1*. Targets of the microRNAs at this locus increase β cell apoptosis^42^, and reduced *Meg3* impairs insulin secretion^43^. *DLK1* inhibits adipocyte differentiation, protecting from obesity^44^, and promotes pancreatic ductal cell differentiation into β cells, increasing insulin secretion^45,46^. Other variants near *MEG3* have been associated with type 1 diabetes (EAS and EUR LD *r*^2^=0 with EAS lead variant)^47^. The other noncoding RNA locus is the *MIR17HG* cluster of miRNAs that regulate glucose-stimulated insulin secretion and pancreatic β cell proliferation stress^48^; one of these microRNAs, miR-19a, affects hepatic gluconeogenesis^49^. Yet another independent T2D association locus is located near *TRAF3*, which is a direct target of the *MIR17HG* microRNA cluster and promotes hyperglycemia by increasing hepatic glucose production^50,51^. The T2D association results suggest that these noncoding RNAs influence disease susceptibility.

## DISCUSSION

These T2D GWAS meta-analyses in the largest number of East Asian individuals analyzed to date identified 56 novel loci, providing additional insight into the biological basis of T2D. The results emphasize substantial shared T2D susceptibility with European individuals, as shown by the strong correlation of effect sizes among T2D-associated genetic variants with common allele frequencies in both East Asian and European ancestry populations. The results also detect novel associations in East Asian individuals, several of which are identified because they have higher allele frequencies in East Asian populations, exhibit larger effect sizes, and/or are influenced by other environmental risk factors or lifestyle behaviors such as alcohol consumption.

The identified loci point to multiple plausible molecular mechanisms and many new candidate genes linking T2D susceptibility to diverse biological processes. Annotation of loci identified in the East Asian meta-analysis suggests a substantial role for insulin resistance in T2D pathogenesis among East Asian individuals through skeletal muscle, adipose, and liver development and function. We also provide evidence that multiple distinct association signals in the same region do not necessarily act through the same gene. Conditionally distinct association signals in close proximity can affect different genes that may act in different tissues by different mechanisms, emphasizing the value of identifying functional variants that enable variant-to-gene links to be examined directly. Our results provide a foundation for future biological research in T2D pathogenesis and offer potential targets for mechanisms for interventions in disease risk.

## METHODS

### Ethics statement

All human research was approved by the relevant institutional review boards for each study at their respective sites and conducted according to the Declaration of Helsinki. All participants provided written informed consent.

### Study cohorts and quality control

The East Asian type 2 diabetes (T2D) meta-analyses were performed with studies participating in the Asian Genetic Epidemiology Network (AGEN), a consortium of genetic epidemiology studies of T2D and related traits conducted in individuals of East Asian ancestry, and the Diabetes Meta-analysis of Trans-ethnic Association Studies (DIAMANTE), a consortium examining the genetic contribution to T2D across diverse ancestry populations including African-American, East Asian, European, Hispanic, and South Asian. The East Asian meta-analysis included 77,418 T2D cases and 356,122 controls from 23 GWAS, including three biobanks, CKB, KBA^52,53^, and BBJ^4^ [effective sample size (N_eff_) = 211,793; Supplementary Figure 1]. A subset of studies was analyzed in BMI-adjusted and sex-specific models (54,481 cases, 224,231 cases; N_eff_ = 135,780). For each study, T2D case control ascertainment is described in Supplementary Table 1 and summary statistics are provided in Supplementary Table 2. Included studies were genotyped on either commercially available or customized Affymetrix or Illumina genome-wide genotyping arrays. Array quality control criteria implemented within each study, including variant call rate and Hardy-Weinberg equilibrium, are summarized in Supplementary Table 3. To harmonize study-level genotype scaffold for imputation to 1000 Genomes (1000G) reference panels, each study adopted a uniform protocol for pre-imputation quality checks. Each study applied the protocol to exclude variants with: i) mismatched chromosomal positions or alleles not present in the reference panel; ii) ambiguous alleles (AT/CG) with minor allele frequency (MAF) >40% in the reference panel; or iii) absolute allele frequency differences > 20% compared to East Asian-specific allele frequencies. The genotype scaffold for each study was then imputed to the 1000G Phase 1 or 3 reference panel^54^ using minimac3^55^ or IMPUTEv2^56^. In BMI-unadjusted analyses, all studies were imputed to 1000G Phase 3. In BMI-adjusted and sex-stratified analyses, all studies were imputed to 1000G Phase 3 except for Biobank Japan^14^, which was imputed to the 1000G Phase 1 reference panel.

### Study-level association analyses

Within each study, all variants were tested for association with T2D assuming an additive model of inheritance within a regression framework, including age, sex, and other study-specific covariates (Supplementary Table 3). To account for population structure and relatedness, association analyses were either performed using FIRTH^57^ or mach2dat with additional adjustment for principal components in unrelated individuals or a linear mixed model with kinship matrix implemented in BOLT-LMM^58^. In studies analyzed with the linear mixed model, allelic effects and standard errors were converted to the log-odds scale that accounts for case-control imbalance^59^. Within each study, variants were removed if the: i) imputation quality score was poor (minimac3 r^2^<0.30; IMPUTE2 info score <0.40); ii) combined case control minor allele count <5; or iii) standard error of the log-OR>10. For a subset of the studies, BMI was added as an additional covariate, and association analyses were also performed separately in males and females. For each study and model, association statistics were corrected with genomic control inflation factor^60^ calculated from common variants (MAF≥5%) (Supplementary Table 3). For BBJ, we applied the genomic control inflation factor 1.21 as reported^4^.

### Sex-combined meta-analysis

We combined study-level association statistics using fixed effects meta-analysis with inverse-variance weighting of log-ORs implemented in METAL^61^. Variants with allele frequency differences >20% between 1000 Genomes Phase 1 and 3 panels were excluded from the meta-analysis. To assess excess inflation arising from cryptic relatedness and population structure, we applied LD score regression to the meta-analysis summary statistics to estimate residual inflation of summary statistics, using a set of 1,889 unrelated Chinese individuals from the Singapore Chinese Eye Study^62^. The LD score regression intercepts were 0.991 for BMI-unadjusted, and 1.0148 for BMI-adjusted models. As the LD score regression intercepts indicated absence of excess inflation, the meta-analysis results were corrected for inflation using these LD score regression intercepts. For subsequent analyses, we considered only variants that were present in at least 50% of the effective sample size N_eff_ [computed as 4/(1/N_cases_ + 1/N_controls_)]^61^. Heterogeneity in allelic effect sizes between studies were assessed with fixed effects inverse variance weighted meta-analysis *P_het_*. We further compared the genetic effects from BMI-unadjusted and BMI-adjusted models using fixed effects inverse variance weighted meta-analysis *P_het_*. Loci were defined as novel if the lead variant is: (1) at least 500 kb away and confirmed by GCTA to be conditionally independent from previously reported T2D-associated variants in any ancestry, and (2) assessed using LocusZoom plots and detailed literature review to be away from known loci with extended LD.

### Sex-differentiated meta-analysis

The meta-analyses described above were repeated for males and females separately. The male-specific meta-analyses included up to 28,027 cases and 89,312 controls (N_eff_ = 65,660) and the female-specific analyses included up to 27,370 cases and 135,055 controls (N_eff_ = 70,332). LD score regression intercepts were 1.0035 for BMI-unadjusted and 1.0034 for BMI-adjusted models in males and 1.0035 for BMI-unadjusted and 1.0034 for BMI-adjusted models in females. We further performed a test for heterogeneity in allelic effects between males and females as implemented in GWAMA^63,64^.

### Detection of distinct association signals

To detect multiple distinct association signals at each associated locus, we combined overlapping loci when the distance between any pair of lead variants was <1 Mb. We then performed approximate conditional analyses using GCTA^8^ with genome-wide meta-analysis summary statistics and LD estimated from 78,000 samples from the Korean Biobank Array^53^.

### Comparing loci effects between East Asian and European populations

We compared the per-allele effect sizes of lead variants identified from the East Asian BMI-unadjusted sex-combined meta-analysis (178 lead variants) and European BMI-unadjusted sex-combined meta-analysis^2^ (231 lead variants; Supplementary Table 7). Across the 409 associated variants from the two ancestries, 11 lead variants overlapped, resulting in 398 unique variants. As the variants in the European analysis were imputed using the Haplotype Reference Consortium reference panel and did not include indel variants, a variant in strong LD (East Asian *r*^2^>0.90) with the lead East Asian variant was used when the lead variant was an indel, when possible. If the lead East Asian variant or a variant in strong LD (East Asian *r*^2^>0.90) was not available in the European data from DIAMANTE, we used results from a previous European type 2 diabetes meta-analysis^65^. The effect size comparison plot was restricted to 343 variants where data was available for both ancestries (Figure 2A). For loci that were significant in both the East Asian and European meta-analyses, if the lead variants were different, both lead variants were plotted (see Supplementary Table 7). Effect size plots were further restricted to: i) 290 lead variants with MAF≥5% in both East Asian and European meta-analyses (Supplementary Figure 7); ii) 162 lead variants significant in the East Asian meta-analysis (Supplementary Figure 8A); and iii) 192 lead variants significant in the European meta-analysis (Supplementary Figure 8B).

### Associations with other metabolic traits and outcomes

We used the Type 2 Diabetes Knowledge Portal^39^ to explore associations of the newly identified loci with other metabolic traits and outcomes. Association statistics from the following consortia were available for query on the portal (last accessed March 18, 2019): coronary artery disease from CARDIoGRAM^66^, BMI and waist-hip-ratio from GIANT^67,68^, lipid traits from GLGC^69^, and glycemic traits from MAGIC^70,71^. Additionally, we used available data from AGEN East Asian meta-analyses for lipids^72^ and adiponectin^73^, along with the phenotypic data from the UK Biobank^74^. Effect sizes were obtained from publicly available summary statistic files.

### Colocalization with expression quantitative trait loci (eQTL)

We searched publicly available eQTL databases such as GTEx^75^ and the Parker lab Islet Browser^17^, to identify *cis*-eQTLs at the novel loci in adipose (subcutaneous and visceral), blood, pancreas, pancreatic islet, and skeletal muscle tissue. We also searched for *cis*-eQTLs in subcutaneous adipose tissue data from the METSIM study^19^. Colocalized eQTLs were identified if the lead expression level-associated variant and the GWAS lead variant were in high LD (*r*^2^>0.80) in Europeans to accommodate the predominantly European eQTL data. Reciprocal conditional analyses were also performed using the METSIM data to determine if the GWAS lead variant and the lead eSNP were part of the same eQTL signal.

### Literature review

We conducted a traditional literature review to identify candidate genes at each novel locus using NCBI Entrez Gene, PubMed and OMIM. We included gene symbols and the following keywords as search terms in PubMed: diabetes, glucose, insulin, islet, adipose, muscle, liver, obesity. A gene was considered a potential candidate if an apparent link to T2D biology existed based on prior studies of gene function.

### Functional annotation and experimentation at *NKX6-1*

We used ENCODE^76^, ChromHMM^77^, and Human Epigenome Atlas^78^ data available through the UCSC Genome Browser to identify candidate variants at the association signal near *NKX6-1* that overlapped open-chromatin peaks, ChromHMM chromatin states, and chromatin-immunoprecipitation sequencing (ChIP-seq) peaks of histone modifications H4K4me1, H3K4me3, and H3K27ac, and transcription factors in the pancreas and pancreatic islets. MIN6 mouse insulinoma cells^79^ and 823/13 rat insulinoma cells^80^ were cultured in DMEM (Sigma) supplemented with 10% FBS, 1mM sodium pyruvate, and 0.1 mM beta-mercaptoethanol. The cell cultures were maintained at 37° C with 5% CO_2_. To measure variant allelic differences in enhancer activity at the *NKX6*-1 locus, we designed oligonucleotide primers (forward: CCCTAGTAATGCCCTTTTTGTT; reverse: TCAGCCTGAGAAGTCTGTGA) with KpnI and Xhol restriction sites, and amplified the 400-bp DNA region (GRCh37/hg19 -chr4: 85,339,430-85,339,829) around rs117624659. As previously described^80^, we ligated amplified DNA from individuals homozygous for each allele into the multiple cloning site of the pGL4.23 (Promega) minimal promoter luciferase reporter vector in both the forward and reverse orientations with respect to the genome. Clones were isolated and sequenced for genotype and fidelity. 2.1×10^5^ MIN6 or 3.0×10^5^ 823/13 cells were seeded per well and grown to 90% confluence in 24-well plates. We co-transfected five independent luciferase constructs and *Renilla* control reporter vector (phRL-TK, Promega) using Lipofectamine 2000 (Life Technologies) and incubated. 48-hours post-transfection, the cells were lysed with Passive Lysis Buffer (Promega). Luciferase activity was measured using the Dual-luciferase Reporter Assay System (Promega) per manufacturer instructions and as previously described^81^.

## ACKNOWLEDGEMENTS

This work was supported by subawards (to X.S and Y.S.C.) from NIDDK U01DK105554 (Jose C. Florez). The authors thank all investigators, staff members and study participants for their contributions to all participating studies. A full list of funding, and individual and study acknowledgements are available in Supplementary Materials.

## AUTHOR CONTRIBUTIONS

Project coordination: K.L.M, X.S. Writing: C.N.S., E.S.T, M.B., M.H., Y.J.K., K.L., K.L.M, X.S. Core analyses: C.N.S., M.H., Y.J.K., K.L., A.K.I., H.J.P., S.M.B., X.S. eQTL lookups: M.vdB, A.L.G. DIAMANTE analysis group: J.E.B., M.B., D.W.B., J.C.C., A.M., M.I.M., M.C.Y.N., L.E.P., W.Zhang., A.P.M. Statistical analysis in individual studies: Y.T., M.H., K.S., X.S., F.T., M.N., C.N.S., K.L., F.B., Y.J.K., S.Moon., C.H.T.T., J.Y., X.G., J.Long., J.F.C., V.J.Y.L., S.H.K., H.S.C., C.H.C. Individual study design and principal investigators: M.Igase., T.Kadowaki., Y.X.W., N.K., M.Y., K.L.M., R.G.W., E.S.T., B.J.K., R.C.W.M., J.I.R., F.M., X.O.S., C.Y.C., W.P.K., T.Y.W., K.S.P., Y.S.C., W.H.H.S., J.Y.W. Genotyping and phenotyping: K.K., A.T., Y.K., T.Y., Y.O., J.B.J., T.Katsuya., M.Isono., S.I., K.Yamamoto., A.H., S.D., W.H., J.S., P.G.L., C.Y., Y.G., Z.B., J.Lv., L.L., Z.C., N.R.L., L.S.A., J.Liu., R.M.vD., S.H., K.Yoon., H.M.J., D.M.S., G.J., A.O.L., B.T., W.Y.S., J.C.N.C., M.Y.H., Y.D.I.C., T.Kawaguchi., Y.B.X., W.Zheng., L.Z., C.C.K., M.A.P., M.G., J.M.Y., C.S., M.L.C., M.S.L., C.M.H., L.M.C., Y.J.H., L.C.C., Y.T.C., F.J.T., J.I.R.

## AUTHOR INFORMATION

Summary-level statistics will be available at the AGEN consortium website https://blog.nus.edu.sg/agen/summary-statistics/ and the Accelerating Medicines Partnership T2D portal http://www.type2diabetesgenetics.org/.

The authors declare no competing interest.

## WEB RESOURCES

Pre-imputation preparation and quality control, http://www.well.ox.ac.uk/~wrayner/tools/

Michigan Imputation Server,https://imputationserver.sph.umich.edu/index.html

EPACTS, https://github.com/statgen/EPACTS

RVTESTS, https://github.com/zhanxw/rvtests

METAL, http://csg.sph.umich.edu/abecasis/metal/

LDSC, https://github.com/bulik/ldsc

GWAMA, https://www.geenivaramu.ee/en/tools/gwama

Type 2 Diabetes Knowledge Portal, http://www.type2diabetesgenetics.org

GTEx Portal, https://gtexportal.org/home/

Parker lab Islet Browser, http://theparkerlab.org/tools/isleteqtl/

---

**Supplementary Figure 1: Flow chart of study design, depicting the different data analyses performed.**

**Supplementary Figure 2: Manhattan plot for East Asian T2D meta-analysis association results in model unadjusted for BMI.** −log_10_(*P*) from fixed effects inverse variance weighted genome-wide meta-analysis association results for each variant (*y*-axis) was plotted against the genomic position (hg19; *x*-axis). Known T2D loci achieving genome-wide significance (*P*<5.0×10^−8^) meta-analysis are shown in blue. Loci achieving genome-wide significance that are previously unreported for T2D association are shown in red.

**Supplementary Figure 3: The relationship between effect size and minor allele frequency.**Odds ratios (*y*-axis) and minor allele frequencies (*x*-axis) for 178 primary association signals from the T2D BMI-unadjusted models.

**Supplementary Figure 4: Regional association plots at seven T2D associated loci with the strongest association *P*-value and more than five distinct association signals in East Asians.**(A) *INS/IGF2*/*KCNQ1*, (B) *CDKN2A*/*B*, (C) *PAX4*/*LEP*, (D) *CDKAL1*, (E) *HHEX*/*IDE*, (F) *CDC123*/*CAMK1D*, and (G) *TCF7L2*. Variants are colored based on East Asian 1000G Phase 3 LD with the lead variants for each association signal, shown as diamonds.

**Supplementary Figure 5: Miami plots of East Asian T2D meta-analysis association results adjusted for BMI.** For each −log_10_(*P*) from fixed effects inverse-variance weighted genome-wide meta-analysis association results for each variant (*y*-axis) was plotted against the genomic position (hg19; *x*-axis). (A) Sex-combined meta-analyses in models unadjusted for BMI (top) and adjusted for BMI (bottom). Both sex-combined models include the same set of studies for comparable sample size. Novel T2D-associated loci are shown in blue (models unadjusted for BMI), purple (models adjusted for BMI), or green (both); (B) sex-specific meta-analyses for males (top) and females (bottom) without BMI adjustment; and (C) sex-specific meta-analyses for males (top) and females (bottom) with BMI adjustment. For (B) and (C), loci significantly associated with T2D in females only are shown in purple and loci significantly associated with T2D in males only are shown in blue.

**Supplementary Figure 6: Effect size comparison of lead variants in sex-combined models unadjusted and adjusted for BMI.** At 178 lead variants identified in the East Asian BMI-unadjusted sex-combined T2D meta-analysis, per-allele effect sizes (β) from BMI-adjusted sex-combined models were plotted against BMI-unadjusted sex-combined model. Both sex-combined models include the same set of studies for comparable sample size. Error bars indicate 95% confidence intervals. Effect sizes between the two models are highly correlated with a Pearson correlation coefficient r=0.99 (Supplementary Table 4).

**Supplementary Figure 7: Regional plots of male-specific T2D-associated locus, *ALDH2*.** (A) Males only, (B) sex-combined, and (C) females only. For each plot, −log_10_(*P*) from association results for each variant (*y*-axis) was plotted against the genomic position (hg19; *x*-axis). The lead variant rs12231737 plotted is the lead variant from BMI-unadjusted male-specific meta-analysis, and also the sex-combined meta-analysis from the same subset of individuals included in the sex-stratified analyses. This lead variant rs12231737 is in high LD with rs77768175, identified from the larger BMI-unadjusted sex-combined meta-analysis (East Asian *r*^2^=0.80). Variants are shaded based on East Asian 1000G Phase 3 LD with the lead variant, shown as a purple diamond.

**Supplementary Figure 8: Effect size comparison of common lead variants (MAF≥5%) identified in East Asian meta-analysis and previously published European T2D GWAS meta-analysis**^2^.

For 290 lead variants with MAF≥5% in both East Asian and European BMI-unadjusted meta-analyses, per-allele effect sizes (β) from Mahajan et al.^2^ (*y*-axis) were plotted against per-allele effect sizes from this East Asian meta-analysis (*x*-axis). Variants are colored purple if they were significant in the East Asian meta-analysis only, green if they were significant in European meta-analysis only, and blue if they were significant in both the East Asian and European meta-analyses. Error bars indicate 95% confidence intervals. (see Methods and Supplementary Table 7).

**Supplementary Figure 9: Effect size comparison of lead variants identified in East Asian BMI-unadjusted meta-analysis and previously published European T2D GWAS meta-analysis**^2^.

For 343 lead variants identified from the two BMI-unadjusted meta-analyses, per-allele effect sizes (β) from a European meta-analysis (*y*-axis) were plotted against per-allele effect sizes from this East Asian meta-analysis (*x*-axis). (A) 162 lead variants significant in the East Asian meta-analysis (purple) or both the East Asian and European meta-analysis (blue) and (B) 192 lead variants significant in the European meta-analysis (green) or both the East Asian and European meta-analysis (blue). These plots include only one variant per locus, in contrast to Figure 2 and Supplementary Figure 8.

**Supplementary Figure 10: T2D-association near *ZNF257* and its relationship with a previously reported *ZNF257* inversion.** The lead T2D-associated variant rs142395395 near *ZNF257* tags an inversion observed almost exclusively in East Asian individuals. We observed an association between the variants tagging the inversion and a decreased risk for T2D. In the reference haplotype (“Reference”), there is no disruption to *ZNF257* and its promoter, thus there is normal ZNF257 function and normal expression of downstream transcripts. When the alternate alleles are present (“Inversion”), an inversion is observed, marked by the separation of the promoter and first two exons of *ZNF257* from the rest of the gene and moving them over 400 kb upstream. While this has not been observed to affect expression of *ZNF208*, *ZNF43*, or *ZNF100,* the inversion results in a loss of ZNF257 function and altered expression of downstream targets^33^.

**Supplementary Figure 11: rs117624659 at *NKX6-1* exhibits allelic differences in transcriptional activity.** (A) Chromatin marks in pancreatic islets in the intergenic region near *NKX6-1* from the UCSC Genome Browser (hg19) spanning the lead T2D associated variant, rs117624659, and the only other candidate variant rs142390274 in high LD (East Asian *r*^2^>0.80) with rs117624659. *NKX6-1* is transcribed from right to left. rs117624659 overlaps regions of accessible chromatin in human pancreatic islets detected by islet FAIRE-seq along with H3K4me1 and H3K4me3 ChIP-seq. It also overlaps a conserved region in vertebrates. The tested candidate regulatory DNA element is represented by a horizontal black rectangle in the upper portion of the figure. (B) rs117624659 at the *NKX6-1* locus exhibited significant allelic differences in transcriptional activity in MIN6 mouse insulinoma cells on a second day. (C-D) rs11762465 at the *NKX6-1* locus did not exhibit significant allelic differences in transcriptional activity in 832/13 rat insulinoma cells on experimental day 1 (C) or day 2 (D).

**Supplementary Figure 12: Forest plots of 49 novel T2D-associated variants using data from five ancestries in the DIAMANTE consortium.** T2D meta-analysis results for the 49 novel T2D-associated variants identified in this East Asian meta-analysis were obtained from the other four ancestry groups within the DIAMANTE consortium (African-American, European, Hispanic, and South Asian) for comparison. Odds ratios (black boxes) and 95% confidence intervals (horizontal lines) for the association between the lead East Asian variants and T2D from each ancestry are presented, along with a combined odds ratio and *P*-value. The size of the box is proportional to the sample size of each contributing ancestry group.

**Supplementary Figure 13: Forest plot of BMI-unadjusted East Asian meta-analysis association results at *SIX3*-*SIX2* locus from each contributing cohort.** Odds ratios (black boxes) and 95% confidence intervals (horizontal lines) for T2D associations at the lead East Asian variant (rs12721928) are presented for each East Asian contributing cohort. The size of the box is proportional to the sample size of each 963 dataset. Full study names can be found in Supplementary Table 1.

