## Supplemental Text for "Identification of type 2 diabetes loci in 433,540 East Asian individuals"

SUPPLEMENTARY MATERIALS

SUPPLEMENTARY MATERIALS	2

Study descriptions	2

Funding, and individual and study acknowledgments	9

REFERENCES	13

### SUPPLEMENTARY MATERIALS

#### Study descriptions

1. The **Anti-Aging Study Cohort (AASC)** includes middle-aged to elderly persons who were consecutive participants in the medical check-up program at the Ehime University Hospital Anti-aging Center^1^. This medical check-up program is provided to general residents of Ehime Prefecture, and is specifically designed to evaluate aging-related disorders, including arteriosclerosis, cardiovascular diseases, physical and cognitive functions. Clinical data used in this study were obtained from the personal medical check-up records of the participants. The ethics committees of Ehime University Graduate School of Medicine approved all study procedures, and written informed consent was obtained from all participants.
2. **Biobank Japan (BBJ)** includes 18,126 subjects enrolled in the BioBank Japan Project at the Institute of Medical Science, the University of Tokyo^2^. Subjects were recruited from 12 medical institutes in Japan including Osaka Medical Center for Cancer and Cardiovascular Diseases, The Cancer Institute Hospital of Japanese Foundation for Cancer Research, Juntendo University, Tokyo Metropolitan Geriatric Hospital, Nippon Medical School, Nihon University School of Medicine, Iwate Medical University, Tokushukai Hospitals, Shiga University of Medical Science, Fukujuji Hospital, National Hospital Organization Osaka National Hospital, and Iizuka Hospital. Subject clinical information including age, gender, height, and weight was collected by a standard questionnaire. All participants provided written informed consent as approved by the ethical committees of the RIKEN Yokohama Institute and the Institute of Medical Science, University of Tokyo. Details of BBJ Type 2 diabetes (T2D) genome-wide association study (GWAS) has been described elsewhere^3^. Data from the Japanese T2D GWAS (2019)^4^ includes subjects from Biobank Japan (BBJ), the Japan Multi-Institutional Collaborative Cohort (J-MICC) Study, the Japan Public Health Center-based Prospective (JPHC) Study, and the Iwate Tohoku Medical Megabank Organization (ToMMo). Details of the study have been described elsewhere^4^.
3. The **Beijing Eye Study (BES)** is a population-based study that assessed the associated and risk factors of ocular and general diseases in a Chinese population^5^. The study was initialized in 2001, collected data from 4439 subjects aged ≥ 40 years from seven communities in Beijing area, where three of the communities were located in rural districts and four were located in urban districts. The BES was followed-up in 2006, with 3251 of the original subjects participating, and in 2011, with 2695 subjects returning for the follow-up examination. At the examinations in 2006 and 2011, fasting blood samples were taken for measurement of blood lipids, glucose, and glycosylated hemoglobin and for genotyping.  All variables used in analyses were taken from examinations in 2006 or in 2011. The BES subjects were genotyped on two arrays, Illumina Human610-Quad (N = 832) and Illumina OmniExpress (N = 814).
4. **Amagasaki Study (CAGE-Amagasaki)** is an ongoing population-based cohort study of 5,745 individuals (3,435 males and 2,310 females), aged >18 years and recruited for a baseline examination between September 2002 to August 2003^6^.
5. **Cardio-metabolic Genome Epidemiology (CAGE-GWAS)** **Network** is an ongoing collaborative effort to investigate genetic and environmental factors, and their interactions affecting cardiometabolic traits/disorders among Asian populations, including the Japanese. CAGE participants were recruited in a population-based or hospital-based setting, depending on the design of member studies^7^. Fasting blood samples were collected after ≥6 hour fast.
6. **Kita-Nagoya Genome Epidemiology Study (CAGE-KING)** is an ongoing community-based prospective observation study of the genetic basis of cardiovascular disease and its risk factors^8^. The study recruited 3,975 Japanese subjects aged 50-80 years who underwent community-based annual health check-ups between May 2005 and December 2007. A subset of the KING study samples were included in the GWAS analysis.
7. **Cebu Longitudinal Health and Nutrition Survey (CLHNS)** is an ongoing community-based birth cohort study that began in 1983. The original study population, study design and recruitment protocols have been described in detail previously^9^. The baseline survey randomly recruited 3,327 pregnant women from the Metropolitan Cebu area, the Philippines in 1983-84 (3,080 singleton live births), and since followed them and their offspring to the present. Trained field staff conducted in-home interviews and collected anthropometric measurements at each visit. Overnight fasting blood samples for biomarker measurement and DNA extraction were obtained in 2005. Details of genotyping has been described elsewhere^10^.
8. **China Health and Nutrition Survey (CHNS)** is a nationwide, longitudinal survey aimed at examining economic, sociological, demographic, and health questions in a Chinese population^11^. Briefly, a stratified probability sample with a multistage, random cluster design was used to select counties and cities within 9 diverse provinces (Guangxi, Guizhou, Heilongjiang, Henan, Hubei, Hunan, Jiangsu, Liaoning, and Shandong), stratified by income and urbanization using State Statistical Bureau definitions. A total of 4,560 households from 228 communities were then randomly selected from within each stratum. Health data was collected during seven rounds of surveys from 1991-2009.
9. **China Kadoorie Biobank (CKB)** is a prospective population-based cohort of 512,891 adults aged 30-79 years recruited from 10 geographically defined regions during 2004-2008^12^. The baseline survey collected questionnaire data, physical measurements, and blood samples, including measurement of blood glucose with recording of time since last meal. Five-yearly resurveys are undertaken among a 5% randomly-selected sample of surviving participants, collecting the same information as at the baseline survey. All participants are followed for cause-specific mortality and morbidity and for any hospital admission, through linkages with registries and health insurance databases. Local, national and international ethics approval was obtained and all participants provided written informed consent.
10. The **Diabetic Cohort (DC)** was initiated in 2004 to investigate the role of genetic, physiological and lifestyle factors in the development of diabetes and its complications^13^. At baseline, >14,000 patients were recruited from primary care facilities in Singapore. Retrospective medical records were obtained five years prior to recruitment. Medical records of >6,000 participants who are being seen at the National Healthcare Group (NHG) polyclinics have been linked to the cohort data up to 2016. From 2011 to 2016, these participants were invited for a first follow-up and a second revisit has been initiated since 2017. Details of T2D GWAS using DC cases and SP2 controls has been described elsewhere^14^.
11. **Singapore Prospective Study Program (SP2)** comprised 6,968 participants in total, of the ages of between 24 and 95 years from four previous cross-sectional studies^15,16^*.* Each of these studies sampled randomly from the Singapore population with a disproportionate sampling scheme to increase the sample sizes of the minority ethnic groups (Malays and Asian Indians). In addition, 10,747 more participants were invited to participate in a follow up study from 2003 to 2007, of which 5,157 people completed the questionnaire and clinical examination.
12. The **Health Examines** (HEXA) cohort is one of the KoGES (The Korean Genome and Epidemiology Study) population-based cohorts which were initiated in 2004 aiming to identify risk factors of life-style related complex diseases such as type 2 diabetes, hypertension, and dyslipidemia^17^. Detailed information of HEXA cohort has been described elsewhere^17^. Briefly, a total of 173,357 participants aged >= 40 years were recruited from the national health examinee registry. As part of Korea Biobank Array (KBA) project, at the end of 2017, about 100K samples were genotyped using KBA^18^. Design, contents, and quality control procedures of KBA have been described elsewhere^18^.
13. The **Cardiovascular Disease Association** (CAVAS) cohort is one of the KoGES population-based cohorts which were initiated in 2004 aiming to identify risk factors of life-style related complex diseases such as type 2 diabetes, hypertension, and dyslipidemia. Detailed information of HEXA cohort has been described elsewhere^17^. Briefly, a total of 28,338 participants aged >= 40 years were recruited from community-dwellers of rural area. As part of Korea Biobank Array project, at the end of 2017, about 20K samples were genotyped using KBA. Design, contents, and quality control procedures of KBA have been described elsewhere^18^.
14. The **Hong Kong T2D Registry (HKDR)** was established in 1995 as a quality improvement program at the Prince of Wales Hospital^19^. Patients referred to the hospital for comprehensive assessment of metabolic control and diabetes complications were enrolled. Referral sources included regional public community clinics, general practitioners, hospital-based clinics, and patients discharged from the Prince of Wales Hospital. Patients were excluded if they were non-Chinese in ethnicity or had type 1 diabetes as defined by acute presentation with diabetic ketoacidosis, heavy ketonuria or continuous requirement of insulin within 1 year of diagnosis, or if unknown diabetes type. The healthy control cohort consisted of 517 subjects from 2 sources: a) 327 healthy adults from a territory-wide health awareness and promotion program who were randomly selected by stratified random sampling with computer-generated codes in accordance to the distribution of occupational groups and hospital staff^20^; and b) 190 individuals without diabetes who were recruited from the community for pharmacogenetics studies in hypertension or hyperlipidaemia^21^.
15. **Korea Association Resource (KARE)** is a part of the Korean Genome Analysis Project (KoGAP) which was initiated in 2007 to discover variants associated with Type 2 Diabetes (T2D) and numerous complex traits. Detailed information has been described elsewhere^22^. Briefly, a total of 10,038 participants aged 40 to 69 were recruited from two population-based cohorts comprising the rural Ansung (n=5,018) and urban Ansan (n=5,020) cohorts. The two KARE cohorts were established as part of the Korean Genome Epidemiology Study (KoGES)^17^. All samples were genotyped with Affymetrix Genome-Wide Human SNP array 5.0. The lipid levels were measured in a fasting state.
16. The **Multi-Ethnic Study of Atherosclerosis (MESA)** is a longitudinal study of the characteristics of subclinical cardiovascular disease (disease detected non-invasively before it has produced clinical signs and symptoms) and the risk factors that predict progression to clinically overt cardiovascular disease or progression of the subclinical disease. MESA researchers study a diverse, population-based sample of 6,814 asymptomatic men and women aged 45-84. Thirty-eight percent of the recruited participants are white, 28 percent African-American, 22 percent Hispanic, and 12 percent Asian, predominantly of Chinese descent. In MESA, fasting blood samples were collected at baseline, processed, and stored at −70°C.
17. The **NAGAHAMA Study** is a community-based longitudinal study in Japan. Participants in the Nagahama study were recruited from general population of Nagahama, a rural city of 125,000 inhabitants located in central Japan (N = 9,764)^23^. Community residents, aged between 30 and 74 years at recruitment, who were living independently without physical impairment or dysfunction, were eligible. All study procedures were approved by the Ethics Committee of Kyoto University Graduate School of Medicine and the Nagahama Municipal Review Board. Written informed consent was obtained from all participants.
18. **Shanghai Breast Cancer Study (SBCS)** included two recruitment phases, as described in detail elsewhere^24,25^. During the initial phase (SBCS-I), 1,459 breast cancer patients and 1,556 controls were recruited between 1996 and 1998 through a rapid case-ascertainment system, and the population-based Shanghai Cancer Registry. Blood samples were obtained from 1,193 (82%) cases and 1,310 (84%) controls. The second phase of participant recruitment (SBCS-II) was conducted between 2002 and 2005 using a protocol similar to the one used during the initial phase. A total of 1,989 incident cases and 1,918 community controls were recruited. The majority of cases (n=1,932, 97.1%) and controls (n=1,857, 96.8%) provided a blood sample or an exfoliated buccal cell sample.
19. **Shanghai Women’s Health Study** (SWHS) is a population-based cohort study of approximately 75,000 women who were aged 40-70 years at study enrollment (1997-2000) and resided in seven geographically defined communities^26^. 56,832 individuals (75.8%) provided a blood sample. A subset of participants who were selected for nested case-control studies of cancer and diabetes with both lipid profile data and GWAS data available were included in this study.
20. **Singapore Chinese Eye Study (SCES)** is a population-based, cross-sectional study to investigate the epidemiology of eye diseases and traits in Singaporean Chinese. Sampling was performed in the 15 residential districts, and included 3,353 Chinese^27^. Details of genotyping in SCES have been described elsewhere^28,29^.
21. **Singapore Chinese Health Study (SCHS)** is a cohort study of 63,257 Singaporean Chinese (Hokkien or Cantonese dialect group) aged 45-74 years, and residing in public housing estates^30,31^. Recruitment and assessment of baseline diet and other interviews took place in the participant’s home from 1993 to 1998. Blood was collected in 28,439 participants between 2000 and 2005. The cohort has been followed up for mortality and morbidity through regular record linkage with the Singapore Cancer Registry, the Hospital Discharge Summary Database and the Singapore Registry of Births and Deaths through collaboration with the Ministry of Health. Details of the T2D GWAS have been described elsewhere^31^.
22. **Singapore Malay Eye Study (SiMES)** is a population-based, cross-sectional study to investigate the epidemiology of eye diseases and traits in Singaporean Malays^32^. 3,280 Malay adults living in 15 residential districts in the southwestern part of Singapore were recruited. Details of Type 2 diabetes (T2D) genome-wide association study (GWAS) has been described elsewhere^14^.
23. **Seoul National University Hospital (SNUH)** study is a hospital based cohort study to investigate the genetic etiology of diabetes and its complications in Koreans. A total of 2,100 T2D cases and 911 controls participated in this study. T2D participants were enrolled from Diabetes Clinic at Seoul National University Hospital and non-diabetic control participants were enrolled from Health Examination Clinic at the same hospital. The details of this study have been described elsewhere^33^.
24. **Samsung Hospital (SSH) study** is a hospital-based study of 331 cases and 118 controls. The age of subjects was mostly ranged between 30-80 years. All individuals provided a blood sample. All DNA samples were genotyped with Affymetrix NSP 250K. SNP imputation was carried out using IMPUTE2 program. The 1000G Phase3 data were used as a reference panel for SNP imputation.
25. The **TAIwanmetaboCHIp consortium (TaiChi)** was formed through a collaborative effort between investigators based in the U.S. and Taiwan. The TaiChi consortium consists of 7 studies that collaborated initially in a large scale metabochip study, and became an ongoing consortium for studies of cardiometabolic disease in the Chinese population in Taiwan. The consortium’s main aim is to identify genetic determinants of atherosclerosis and diabetes related traits in East Asians, and to fine map validated loci identified in other race/ethnic groups. The main academic sites in Taiwan include Taipei and Taichung Veteran’s General Hospitals (VGH), National Health Research Institute (NHRI), Tri-Service General Hospital (TSGH), and National Taiwan University Hospital (NTUH). The main academic sites participating in the TaiChi consortium from United States include Stanford University School of Medicine in Stanford, California; Hudson-Alpha Biotechnology Institute in Huntsville, Alabama; and Los Angeles Biomedical Research Institute at Harbor‑UCLA Medical Center in Torrance California. There are 7 principal cohorts that consisted of in total 11,859 samples. The detailed description of each cohort has been published elsewhere^34^. The seven studies included the following: 1) Taiwan Diabetes and RelAted Genetic COmplicatioN (**Taiwan DRAGON**), a cohort study of type 2 diabetes at Taichung Veterans General Hospital (Taichung VGH) in Taiwan, with participants including individuals with either newly diagnosed or established diabetes (subjects with hyperglycemia who did not meet diagnostic criteria for type 2 diabetes were not included); 2) Healthy Aging Longitudinal Study in Taiwan (**HALST**), a population based epidemiologic study of older adults living in all major geographic regions of Taiwan established by the Taiwan National Health Research Institutes (NHRI); 3) Stanford-Asian Pacific Program in Hypertension and Insulin Resistance (**SAPPHIRe**), a family based study established in 1995 with an initial goal of identifying major genetic loci underlying hypertension and insulin resistance in East Asian populations, with Taiwanese subjects participating in the TaiChi consortium; 4) TAiwan Coronary and Transcatheter intervention (**TACT**), a cohort study that enrolled patients with angina pectoris and objective documentation of myocardial ischemia who underwent diagnostic coronary angiography and/or revascularization any time after October 2000 at the National Taiwan University Hospital (NTUH); 5) Taichung CAD study (**TCAD**), includes patients with a variety of cardiovascular diseases who received care at the Taichung Veterans General Hospital (Taichung VGH), i.e. specifically individuals who were hospitalized for diagnostic and interventional coronary angiography examinations and treatment; 6) Taiwan Coronary Artery Disease GENetic (**TCAGEN**), a cohort study that enrolled patients undergoing coronary angiography or percutaneous intervention at the National Taiwan University Hospital (NTUH) in the setting of either stable angina pectoris or prior myocardial infarction; 7) Taiwan US Diabetic Retinopathy (**TUDR**) enrolled subjects with type 2 diabetes who received care at Taichung Veteran General Hospital (Taichung VGH), and a small number of subjects from Taipei Tri-Service General Hospital (TSGH); TUDR subjects underwent a complete ophthalmic and fundus examination to carefully document the presence and extent of retinopathy. From these 7 studies, samples for over 1,800 subjects were selected based on completeness of standard metabolic phenotyping and knowledge of cardiac disease status, to undergo GWAS genotyping with the Zhonghua chip (**TaiChi-G**).
26. The **Taiwan T2D Study (TWT2D)** includes a total of 2,798 unrelated individuals with T2D, age > 20 years, recruited from China Medical University Hospital (CMUH), Taichung, Taiwan; Chia-Yi Christian Hospital (CYCH), Chia-Yi, Taiwan; and National Taiwan University Hospital (NTUH), Taipei, Taiwan. The controls were randomly selected from the Taiwan Han Chinese Cell and Genome Bank. Details of T2D GWAS has been described elsewhere^35^.

#### Funding, and individual and study acknowledgments

| **Study acronym** | **Study Funding/Acknowledgements** |
| --- | --- |
| AASC | The **Anti-Aging Study Cohort (**AASC) was supported by the Grant-in-Aid for Scientific Research (20018020, 19659163, 20390185, 23659382, 24390084, 23659352, 25293141, 26670313, 17H04123) from Ministry of Education, Culture, Sports, Science and Technology of Japan, research grant from the Japan Atherosclerosis Prevention Found, National Cardiovascular Research Grants, and Research Promotion Award from Ehime University. |
| BBJ | We acknowledge all the staff in the BioBank Japan (BBJ) project as well as the doctors and staff of the contributing hospitals for their outstanding work on collecting samples and clinical information. We also would like to thank all the patients participating in this project. This research was supported by the Tailor-Made Medical Treatment Program (the BioBank Japan Project) of the Ministry of Education, Culture, Sports, Science, and Technology (MEXT) and the Japan Agency for Medical Research and Development (AMED). |
| BES | The Beijing Eye Study (BES) was supported by National Natural Science Foundation of China (grant 81570835). |
| CAGE-Amagaski | We thank Drs. Toshio Ogihara, Yukio Yamori, Akihiro Fujioka, Chikanori Makibayashi, Sekiharu Katsuya, Ken Sugimoto, Kei Kamide, and Ryuichi Morishita and the many physicians of the participating hospitals and medical institutions in Amagasaki Medical Association for their assistance in collecting the DNA samples and accompanying clinical information. |
| CAGE-GWAS | The CAGE Network studies were supported by grants for the Core Research for Evolutional Science and Technology (CREST) from the Japan Science Technology Agency; KAKENHI (Grant-in-Aid for Scientific Research) from the Ministry of Education, Culture, Sports, Science and Technology of Japan; and the Grant and research budget of National Center for Global Health and Medicine (NCGM). |
| CAGE-KING | The KING Study was supported in part by Grants-in-Aid from MEXT (nos. 24390169, 16H05250, 15K19242, 16H06277) as well as by a grant from the Funding Program for Next-Generation World-Leading Researchers (NEXT Program, no. LS056). |
| CHNS | The China Health and Nutrition Survey (CHNS) was supported by the China National Institute for Nutrition and Health; the Chinese Center for Disease Control and Prevention; the National Institutes of Health (R01HD30880, R01HL108427, and R01DK104371); the Fogarty International Center of the National Institutes of Health (TW009077); the China-Japan Friendship Hospital; the Chinese Ministry of Health; the Chinese National Human Genome Center at Shanghai; and the Carolina Population Center (P2CHD050924). |
| CKB | The China Kadoorie Biobank (CKB) chief acknowledgment is to the participants, project staff, and China National Centre for Disease Control and Prevention (CDC) and its regional offices. China’s National Health Insurance provides electronic linkage to all hospital treatment. Funding sources: Baseline survey and first re-survey: Hong Kong Kadoorie Charitable Foundation; long-term follow-up: UK Wellcome Trust (202922/Z/16/Z, 104085/Z/14/Z, 088158/Z/09/Z), National Natural Science Foundation of China (81390540, 81390541, 81390544), and National Key Research and Development Program of China (2016YFC 0900500, 0900501, 0900504, 1303904). DNA extraction and genotyping supported by GlaxoSmithKline and the UK Medical Research Council (MC-PC-13049, MC-PC-14135). The project was support by British Heart Foundation, UK MRC and Cancer Research UK through core funding to the Clinical Trial Service Unit and Epidemiological Studies Unit at Oxford University. |
| CLHNS | The Cebu Longitudinal Health and Nutrition Survey (CLHNS) was supported by US National Institutes of Health grants DK078150, TW005596 and HL085144; pilot funds from RR020649, ES010126, and DK056350; and the Office of Population Studies Foundation. |
| DC/SP2 | The Diabetic Cohort (DC) and Singapore Prospective Study Program (SP2) was supported by the individual research grant and clinician scientist award schemes from the National Medical Research Council (NMRC) and the Biomedical Research Council (BMRC) of Singapore, and NMRC LCG SG100K. |
| HEXA+CAVAS | The Korean Biobank Array (KBA) was supported by grants from Korea Centers for Disease Control and Prevention (4845–301, 4851–302, 4851–307) and intramural grants from the Korea National Institute of Health (2016-NI73001-00, 2019-NG-053-00). This study was performed with bioresources from National Biobank of Korea, the Centers for Disease Control and Prevention, Republic of Korea. Genotype data were provided by the Collaborative Genome Program for Fostering New Post-Genome Industry (3000-3031b). |
| HKDR | The Hong Kong Diabetes Registry (HKDR) acknowledge support from the Theme-based Research Scheme from the Research Grants Council of the Hong Kong Special Administrative Region, China (Project no: T12-402/13-N), the Hong Kong Foundation for Research and Development in Diabetes, the Vice-Chancellor One-off Discretionary Fund, the Focused Innovations Scheme and the Postdoctoral Fellowship Scheme of the Chinese University of Hong Kong. |
| KARE | The Korean Association Resource (KARE) was supported by grants from Korea Centers for Disease Control and Prevention (4845–301, 4851–302, 4851–307) and intramural grants from the Korea National Institute of Health (2016-NI73001-00, 2019-NG-053-00). This study was performed with bioresources from National Biobank of Korea, the Centers for Disease Control and Prevention, Republic of Korea. |
| MESA | MESA and the MESA SHARe project are conducted and supported by the US National Heart, Lung, and Blood Institute (NHLBI) in collaboration with MESA investigators. Support for MESA is provided by contracts HHSN268201500003I, N01-HC-95159, N01-HC-95160, N01-HC-95161, N01-HC-95162, N01-HC-95163, N01-HC-95164, N01-HC-95165, N01-HC-95166, N01-HC-95167, N01-HC-95168, N01-HC-95169, UL1-TR-000040, UL1-TR-001079, UL1-TR-001420. The provision of genotyping data was supported in part by the National Center for Advancing Translational Sciences, TSCI grant UL1TR001881, and the National Institute of Diabetes and Digestive and Kidney Disease Diabetes Research (DRC) grant DK063491. Funding for SHARe genotyping was provided by NHLBI Contract N02-HL-64278.  Genotyping was performed at Affymetrix (Santa Clara, California, USA) and the Broad Institute of Harvard and MIT (Boston, Massachusetts, USA) using the Affymetrix Genome-Wide Human SNP Array 6.0. |
| NAGAHAMA | The **Nagahama** study supported by a university grant, The Center of Innovation Program, The Global University Project, and a Grant-in-Aid for Scientific Research (25293141, 26670313, 26293198, 17H04182, 17H04126, 17H04123, 18K18450) from the Ministry of Education, Culture, Sports, Science and Technology of Japan; the Practical Research Project for Rare/Intractable Diseases (ek0109070, ek0109070, ek0109196, ek0109348), the Comprehensive Research on Aging and Health Science Research Grants for Dementia R&D (dk0207006, dk0207027), the Program for an Integrated Database of Clinical and Genomic Information (kk0205008), the Practical Research Project for Life-style-related Diseases including Cardiovascular Diseases and Diabetes Mellitus (ek0210066, ek0210096, ek0210116), and the Research Program for Health Behavior Modification by Utilizing IoT (le0110005) from Japan Agency for Medical Research and Development (AMED); Welfare Sciences Research Grants, Research on Region Medical from the Ministry of Health, Labor and Welfare of Japan; the Takeda Medical Research Foundation, and the Mitsubishi Foundation, Daiwa Securities Health Foundation, and Sumitomo Foundation. |
| SBC/SWHS | The Shanghai Breast Cancer Study (SBCS) was supported in part by US National Institutes of Health grants R01CA64277 and R01CA124558 , as well as Ingram Professorship and Research Reward funds from the Vanderbilt University School of Medicine. We want to thank participants and research staff of the SBCS, Ms. Regina Courtney for plasma and DNA sample preparation, Drs. Hui Cai, Ben Zhang and Ms. Jing He for data processing and analyses. |
| SCES | The Singapore Chinese Eye Study (SCES), and Singapore Malay Eye Study (SiMES) are supported by the National Medical Research Council (NMRC), Singapore (grants 0796/2003, 1176/2008, 1149/2008, STaR/0003/2008, 1249/2010, CG/SERI/2010, CIRG/1371/2013, and CIRG/1417/2015), and Biomedical Research Council (BMRC), Singapore (08/1/35/19/550 and 09/1/35/19/616). |
| SCHS | The Singapore Chinese Health Study (SCHS) was supported by the US National Institutes of Health grants R01DK08072, R01CA144034 and UM1CA182876. |
| SiMES | The Singapore Chinese Eye Study (SCES), and Singapore Malay Eye Study (SiMES) are supported by the National Medical Research Council (NMRC), Singapore (grants 0796/2003, 1176/2008, 1149/2008, STaR/0003/2008, 1249/2010, CG/SERI/2010, CIRG/1371/2013, and CIRG/1417/2015), and Biomedical Research Council (BMRC), Singapore (08/1/35/19/550 and 09/1/35/19/616). |
| SNUH | This research conducted at the Seoul National University Hospital (SNUH) was supported by a grant from the Korea Health Technology R&D Project through the Korea Health Industry Development Institute, funded by the Ministry of Health & Welfare (grant number: HI15C1595, HI14C0060, HI15C3131). |
| SSH | The Samsung Hospital study (SSH) was supported by the National Research Foundation of Korea (NRF) Grant funded by the Ministry of Science and ICT (NRF-2017R1A2B4006508) |
| TaiChi-G | The TAICHI-G study was supported by grants from the National Health Research Institutes, Taiwan (PH-099-PP-03, PH-100-PP-03, and PH-101-PP-03); the National Science Council, Taiwan (NSC 101-2314-B-075A-006-MY3, MOST 104-2314-B-075A-006-MY3, MOST 104-2314-B-075A-007, and MOST 105-2314-B-075A-003); and the Taichung Veterans General Hospital, Taiwan (TCVGH-1020101C, TCVGH-1020102D, TCVGH-1023102B, TCVGH-1023107D, TCVGH-1030101C, TCVGH- 1030105D, TCVGH-1033503C, TCVGH-1033102B, TCVGH-1033108D, TCVGH-1040101C, TCVGH-1040102D, TCVGH-1043504C, and TCVGH-1043104B); it was also supported in part by the National Center for Advancing Translational Sciences (CTSI grant UL1TR001881). |
| TWT2D | The Taiwan T2D (TWT2D) study was supported by GMM Study, Academia Sinica, Taiwan. |
| **Individual** | **Funding/Acknowledgements** |
| Jennifer E Below | US National Institutes of Health R01AG061351, 5U01DK105554 |
| Donald W Bowden | US National Institutes of Health U01DK105556 and R01DK66358 |
| Fiona Bragg | BHF Centre of Research Excellence, Oxford (RE/13/1/30181) |
| Sarah M Brotman | US National Institutes of Health T32GM007092 |
| John C Chambers | Singapore Ministry of Health’s National Medical Research Council under its Singapore Translational Research Investigator (STaR) Award (NMRC/STaR/0028/2017); iHealth-T2D, 643774; The work was carried out in part at the National Institute for Health Research/Wellcome Trust Imperial Clinical Research Facility. |
| Yoon Shin Cho | National Research Foundation of Korea (NRF) Grant funded by the Ministry of Science and ICT (NRF-2017R1A2B4006508) |
| Ching-Yu Cheng | National Medical Research Council (NMRC), Singapore (CSA-SI/0012/2017) |
| Shufa Du | US National Institutes of Health Fogarty grant D43 TW009077 |
| Anna L Gloyn | ALG is a Wellcome Senior Fellow in Basic Biomedical Science. This work was funded in Oxford by the Wellcome Trust (095101, 200837, 106130, 203141). This work was also supported by the Oxford NIHR Biomedical Research Centre. The views expressed in this article are those of the author(s) and not necessarily those of the NHS, the NIHR, or the Department of Health. |
| Soo-Heon Kwak | Korea Health Technology R&D Project through the Korea Health Industry Development Institute (grant number: HI15C3131) |
| Nannette R Lee | US National Institutes of Health TW008288 |
| Ronald CW Ma | Research Grants Council Theme-based Research Scheme (T12-402/13-N) |
| Karen L Mohlke | US National Institutes of Health R01DK072193, R01DK093757, U01DK105561 |
| Maggie CY Ng | US National Institutes of Health U01DK105556 and R01DK66358 |
| Kyong-Soo Park | Korea Health Technology R&D Project through the Korea Health Industry Development Institute (grant number: HI15C1595, HI14C0060) |
| Hannah J Perrin | US National Institutes of Health T32HL069768 |
| Lauren E Petty | US National Institutes of Health R01AG061351, 5U01DK105554 |
| Andrew P Morris | US National Institutes of Health U01DK105535 |
| Cassandra N Spracklen | American Heart Association Postdoctoral Fellowship 15POST24470131 and 17POST33650016 |
| Martijn van de Bunt | M.vdB. was supported by a Novo Nordisk postdoctoral fellowship run in partnership with the University of Oxford, and currently a full-time employee of Novo Nordisk |
| Jer-Yuarn Wu | Academia Sinica GMM Study |
| Weihua Zhang | iHealth-T2D, 643774; The work was carried out in part at the National Institute for Health Research/Wellcome Trust Imperial Clinical Research Facility. |
