## Supplemental Figures for "Identification of type 2 diabetes loci in 433,540 East Asian individuals"

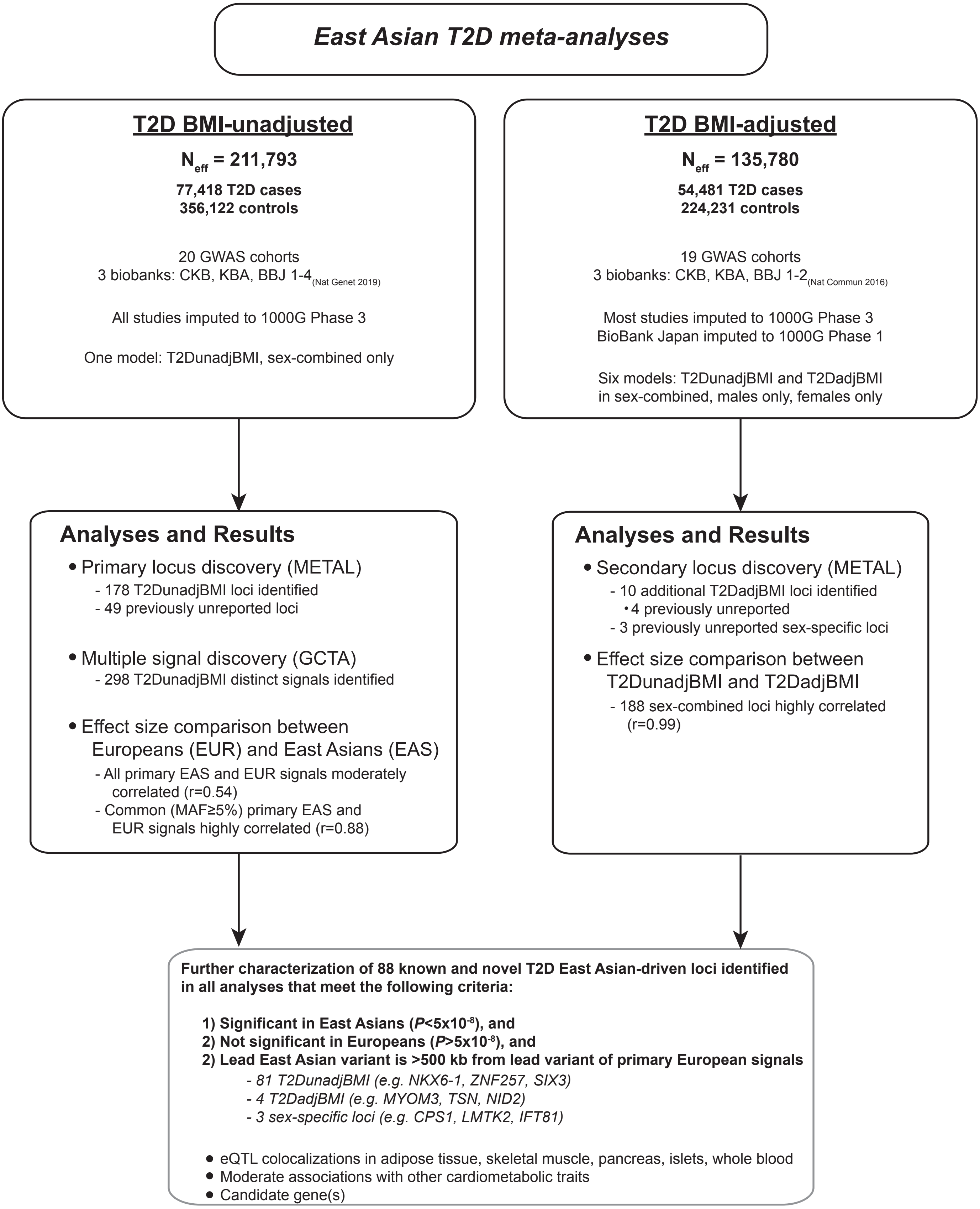


Supplementary Figure 1

Supplementary Figure 2

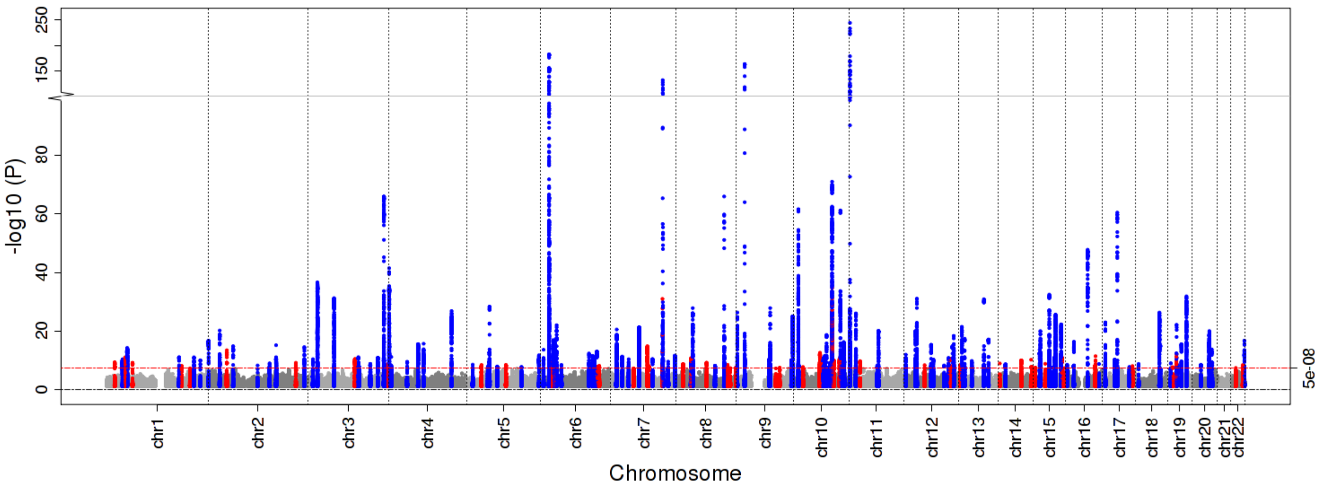

Supplementary Figure 3

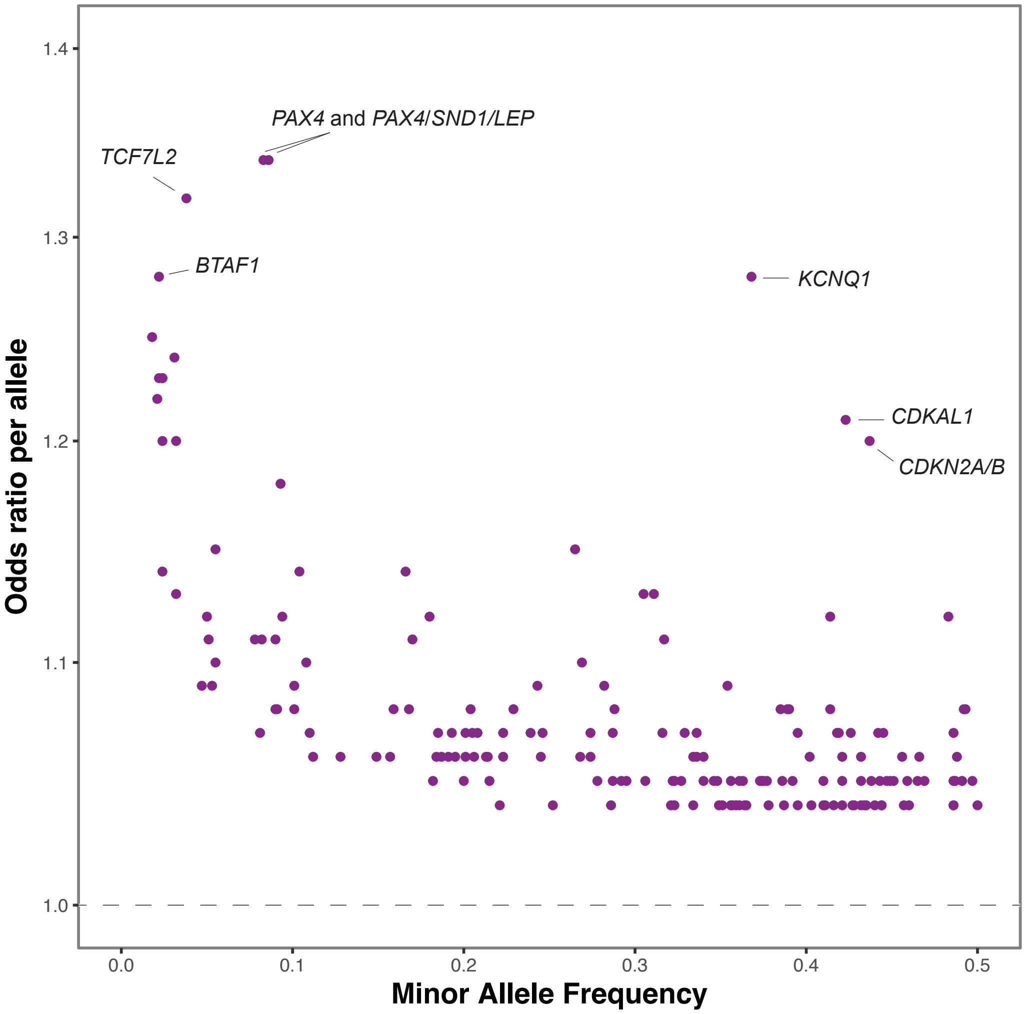

Supplementary Figure 4

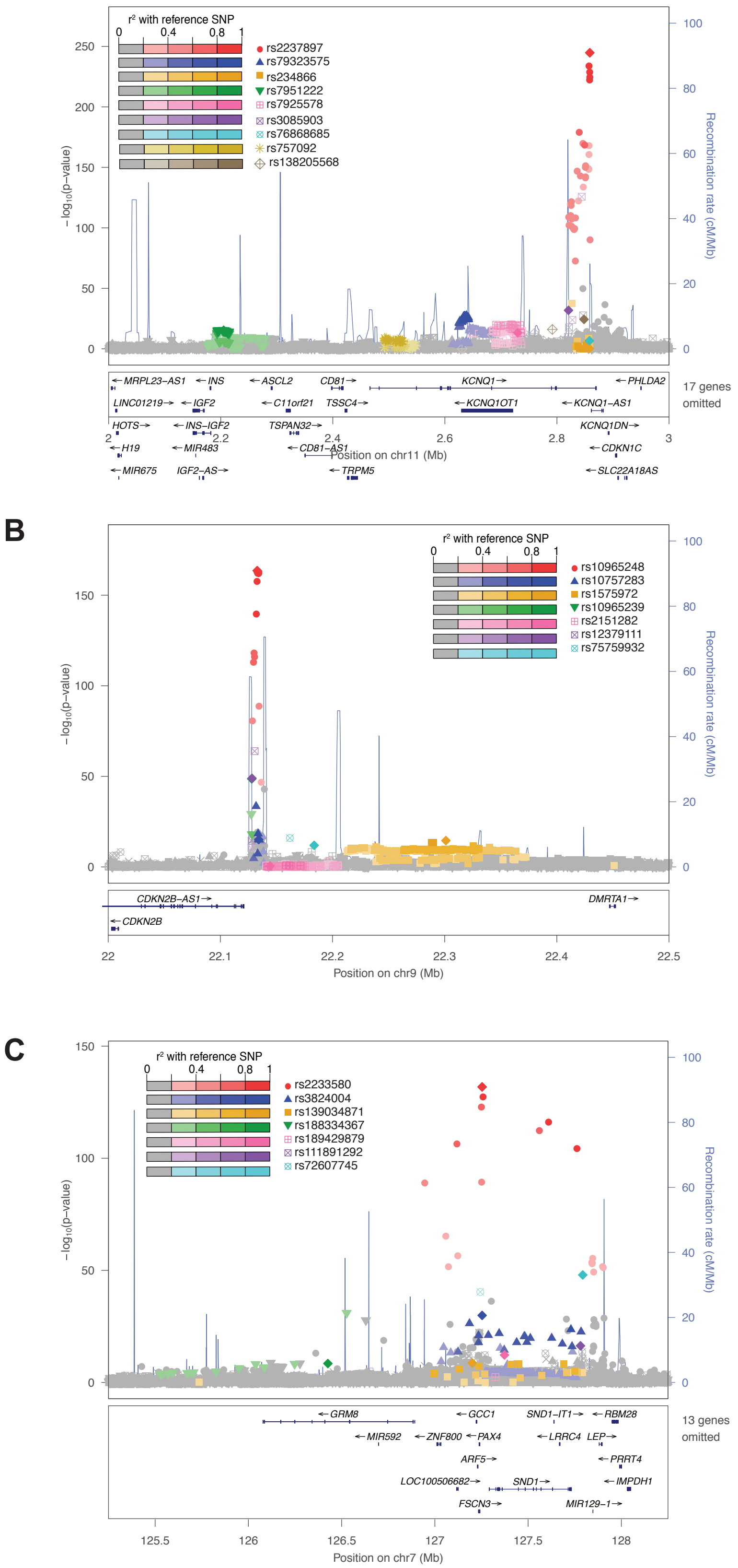

Supplementary Figure 4, continued

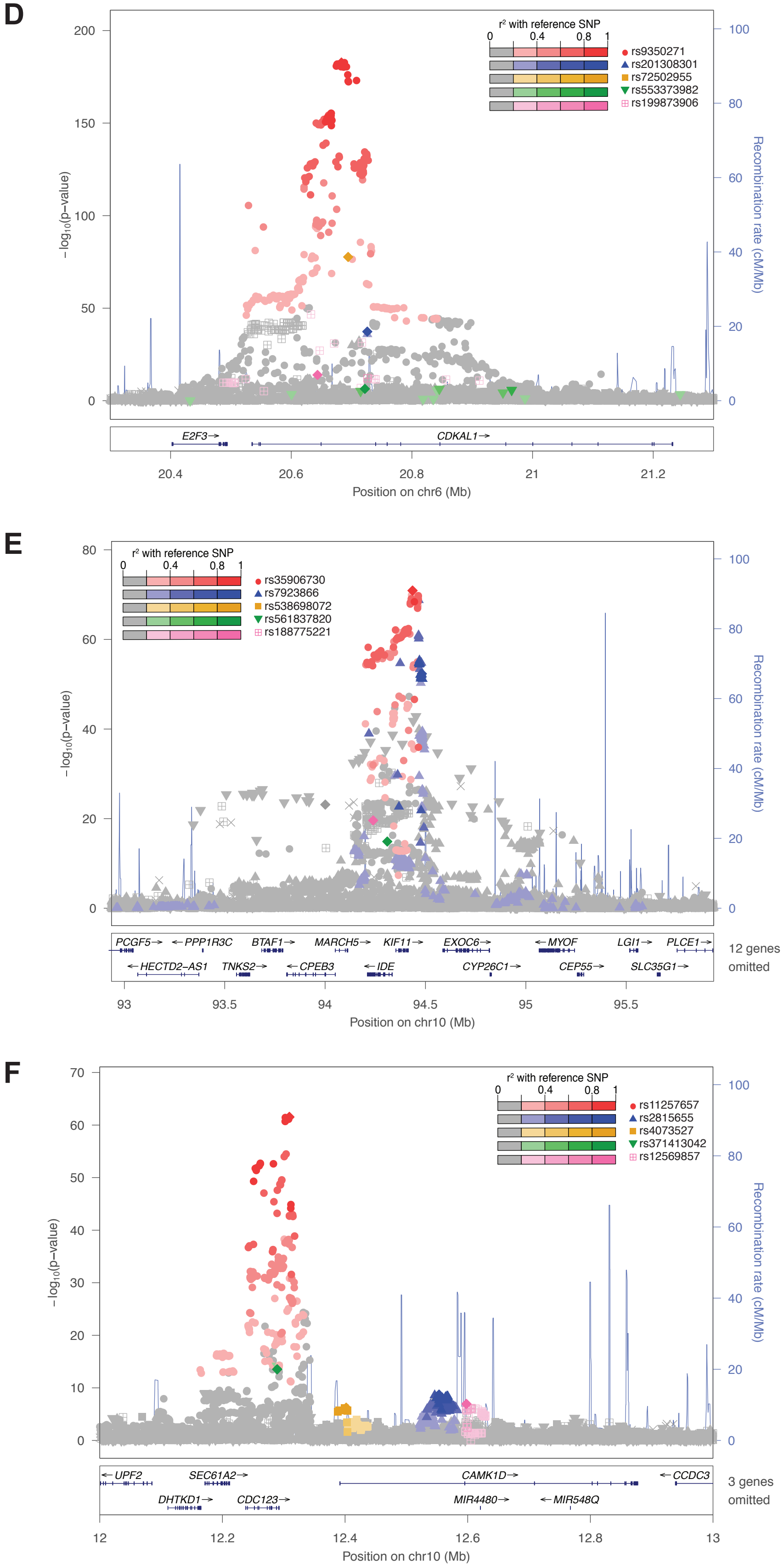

G

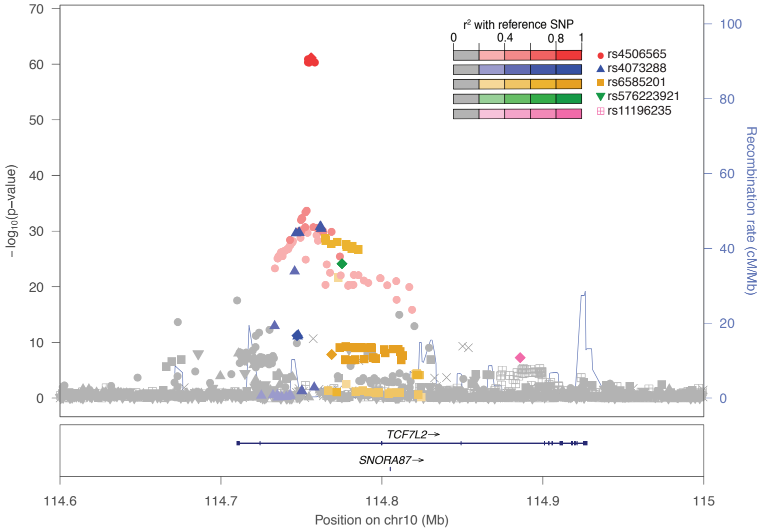


Supplementary Figure 4, continued

Supplementary Figure 5

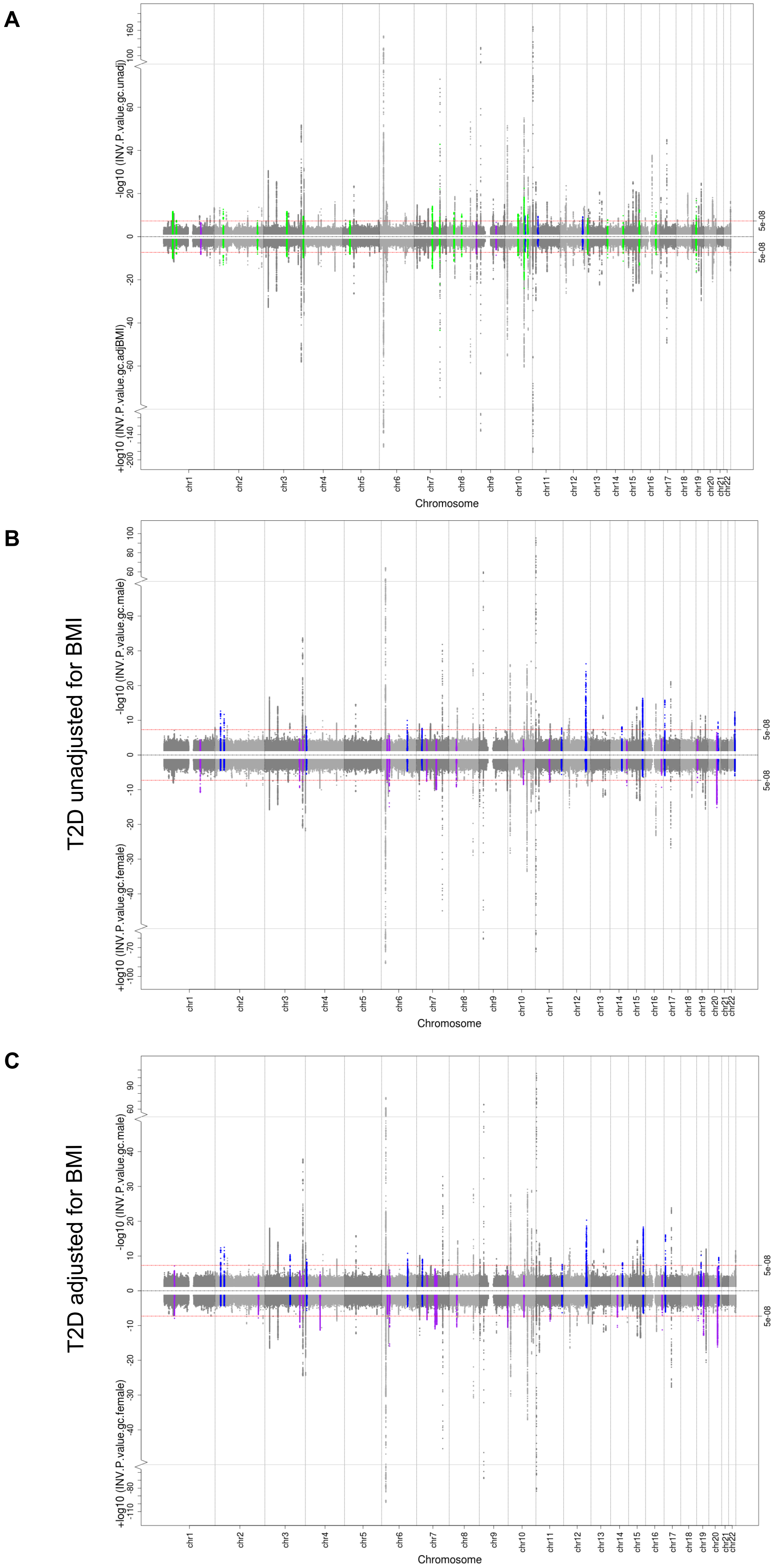

Supplementary Figure 6

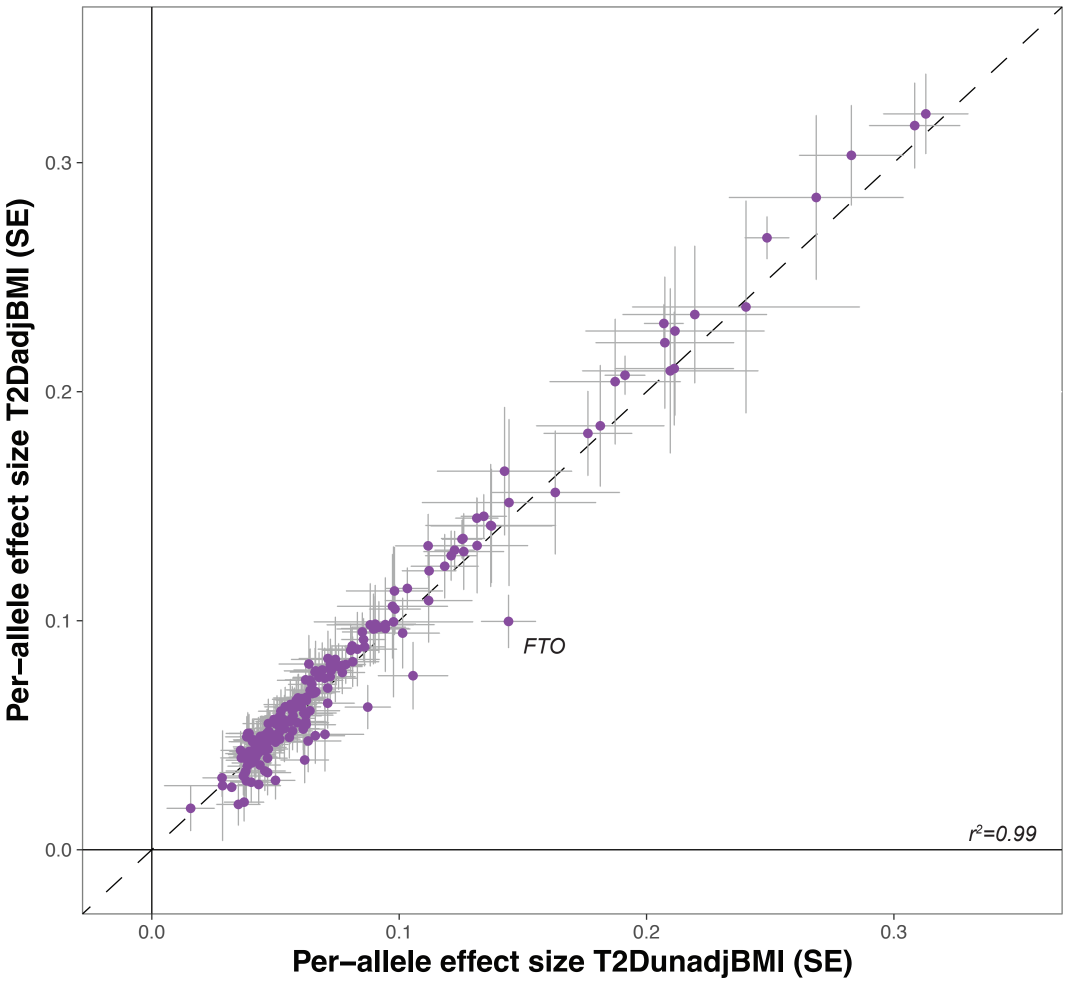

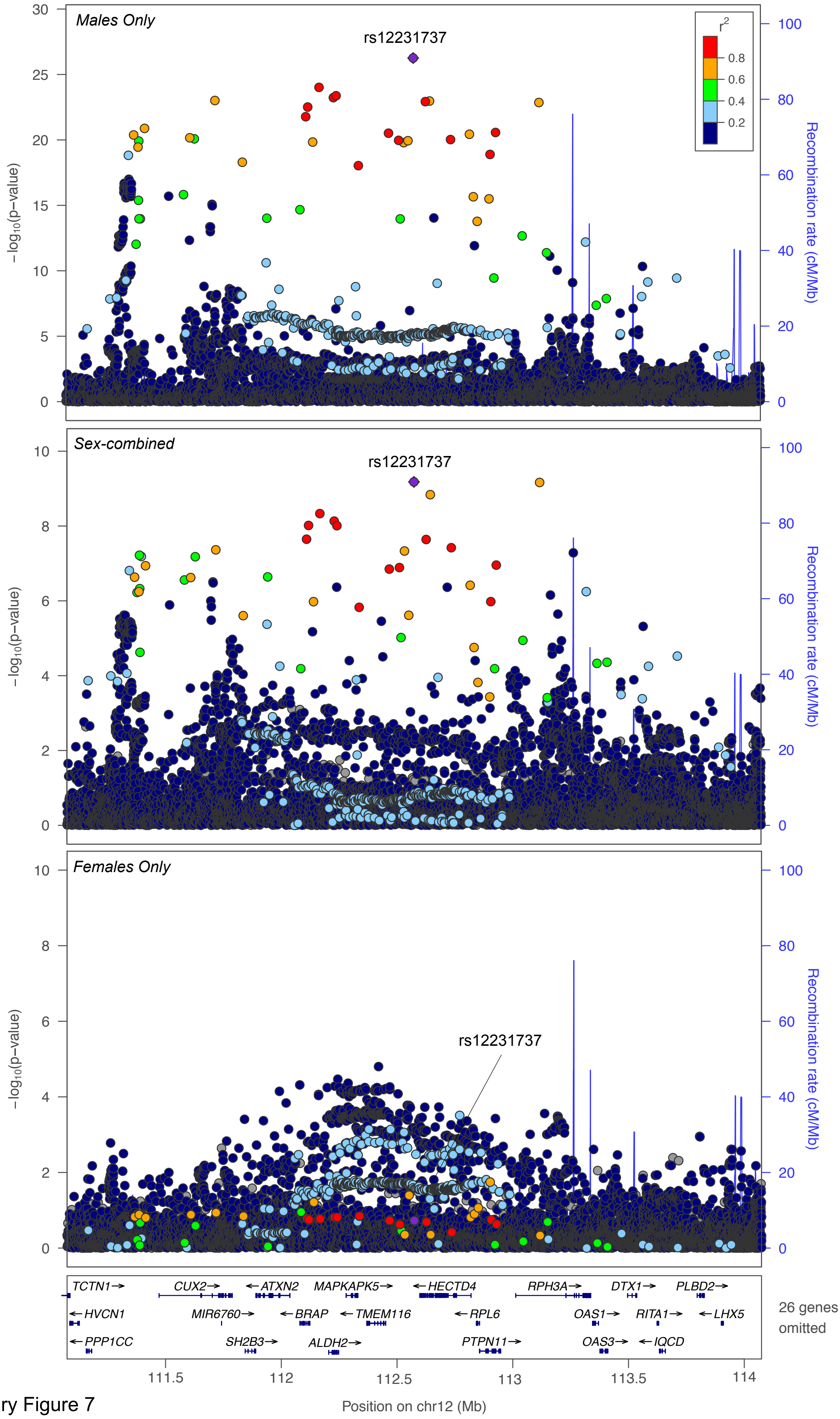


Supplementary Figure 7

Supplementary Figure 8

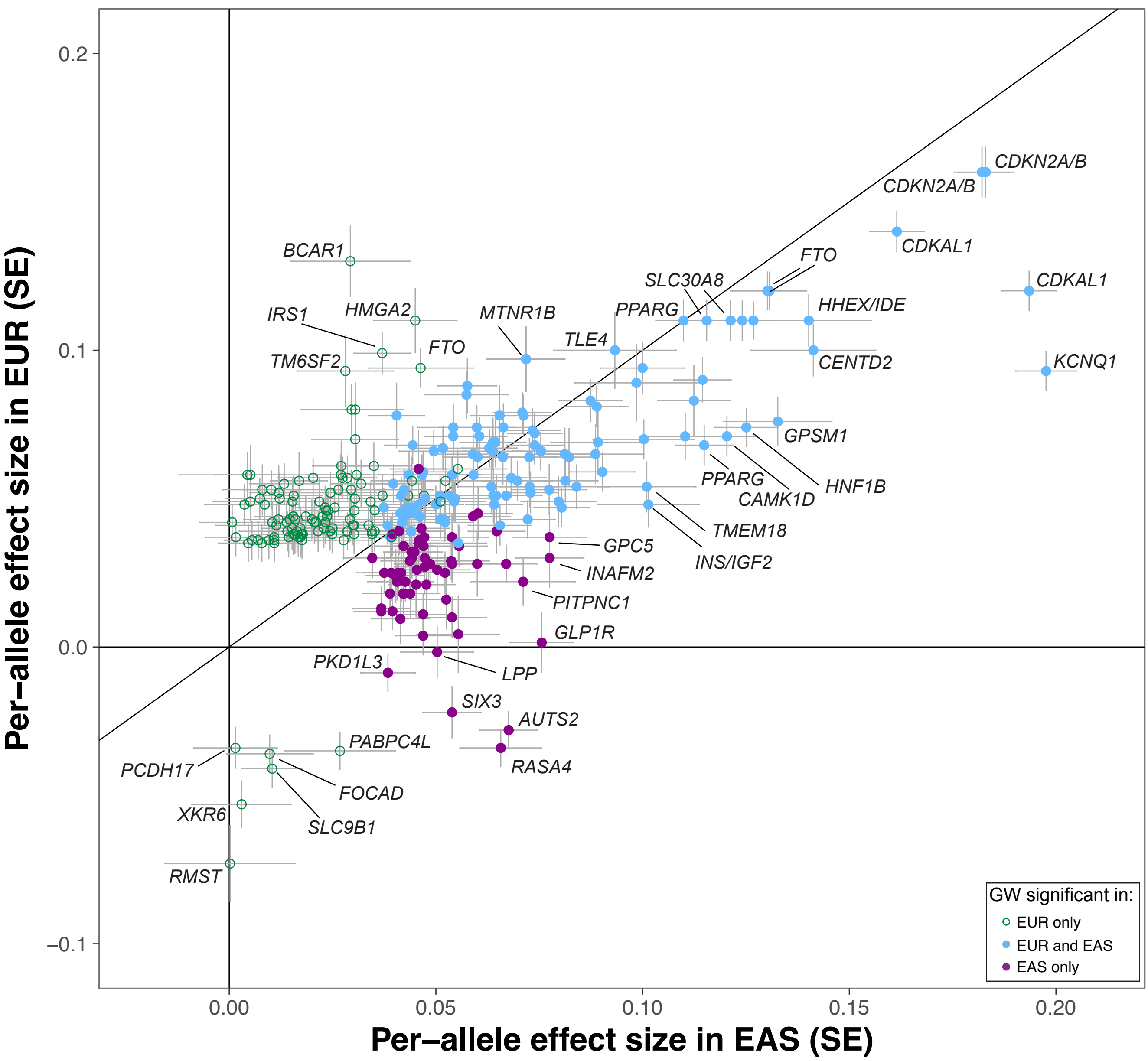

Supplementary Figure 9

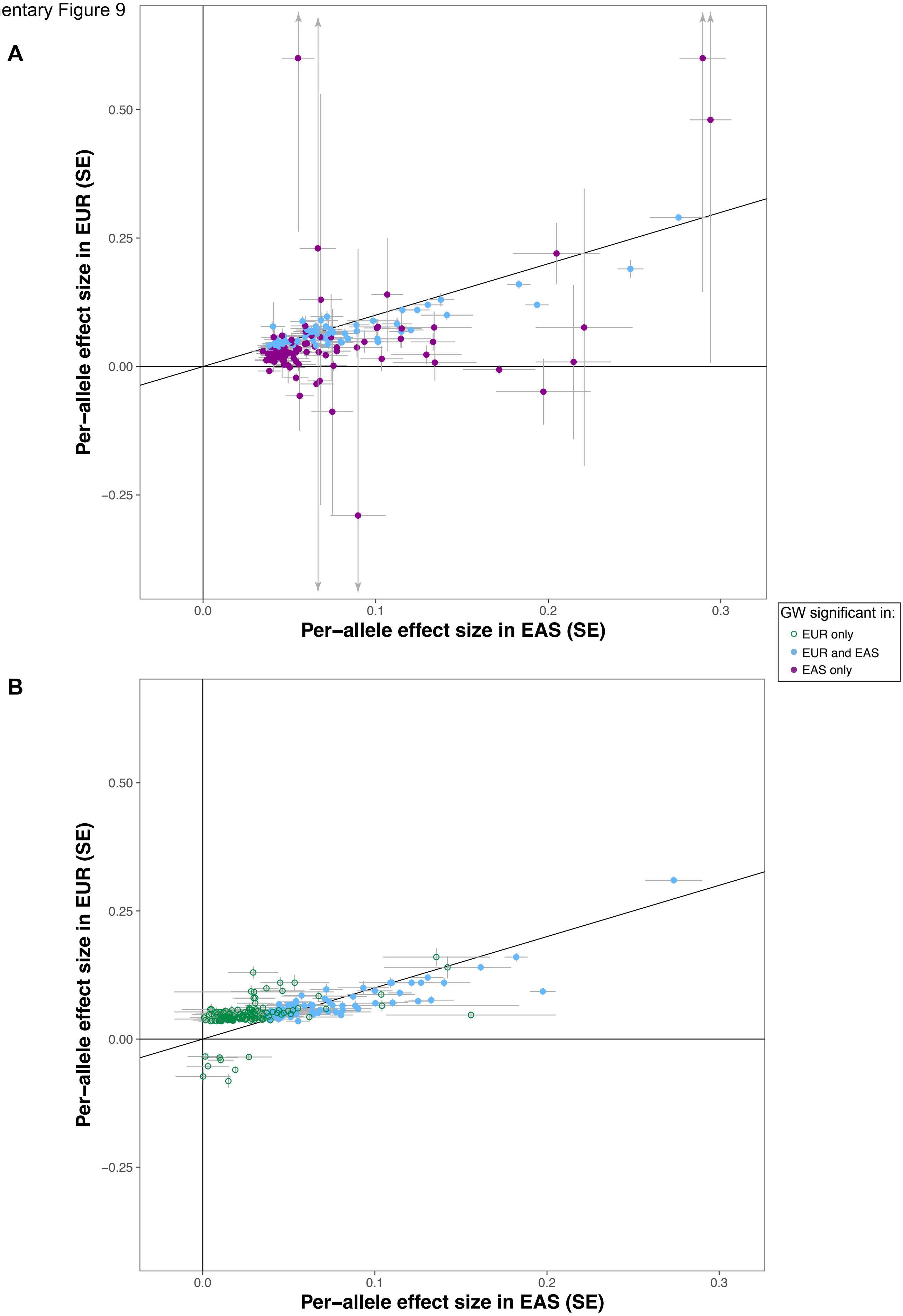

#### Supplementary Figure 10

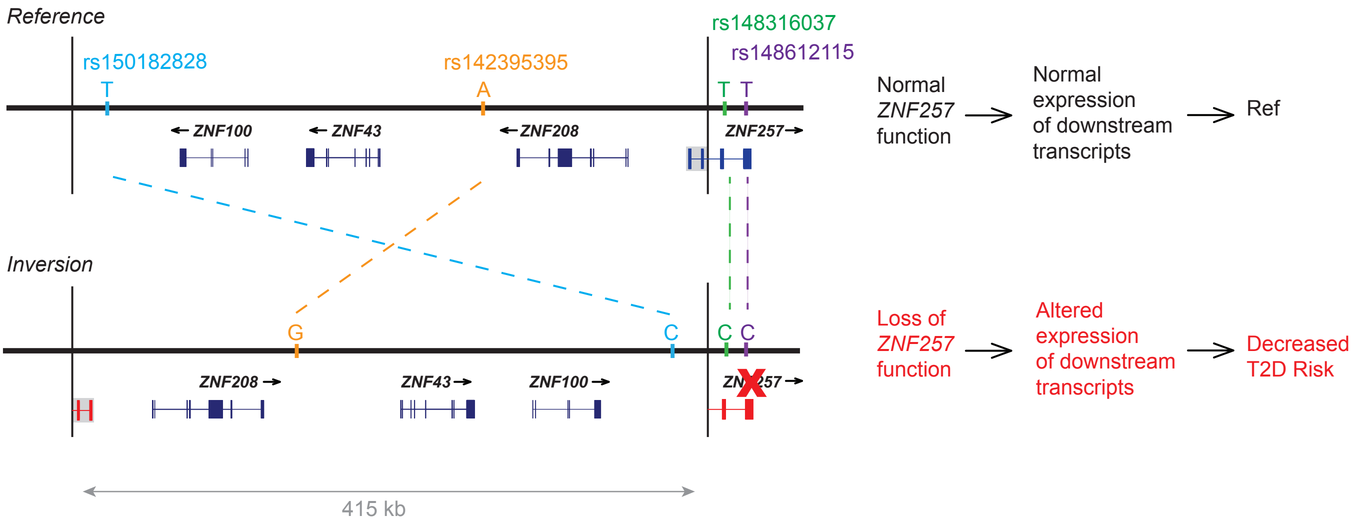

Supplementary Figure 11

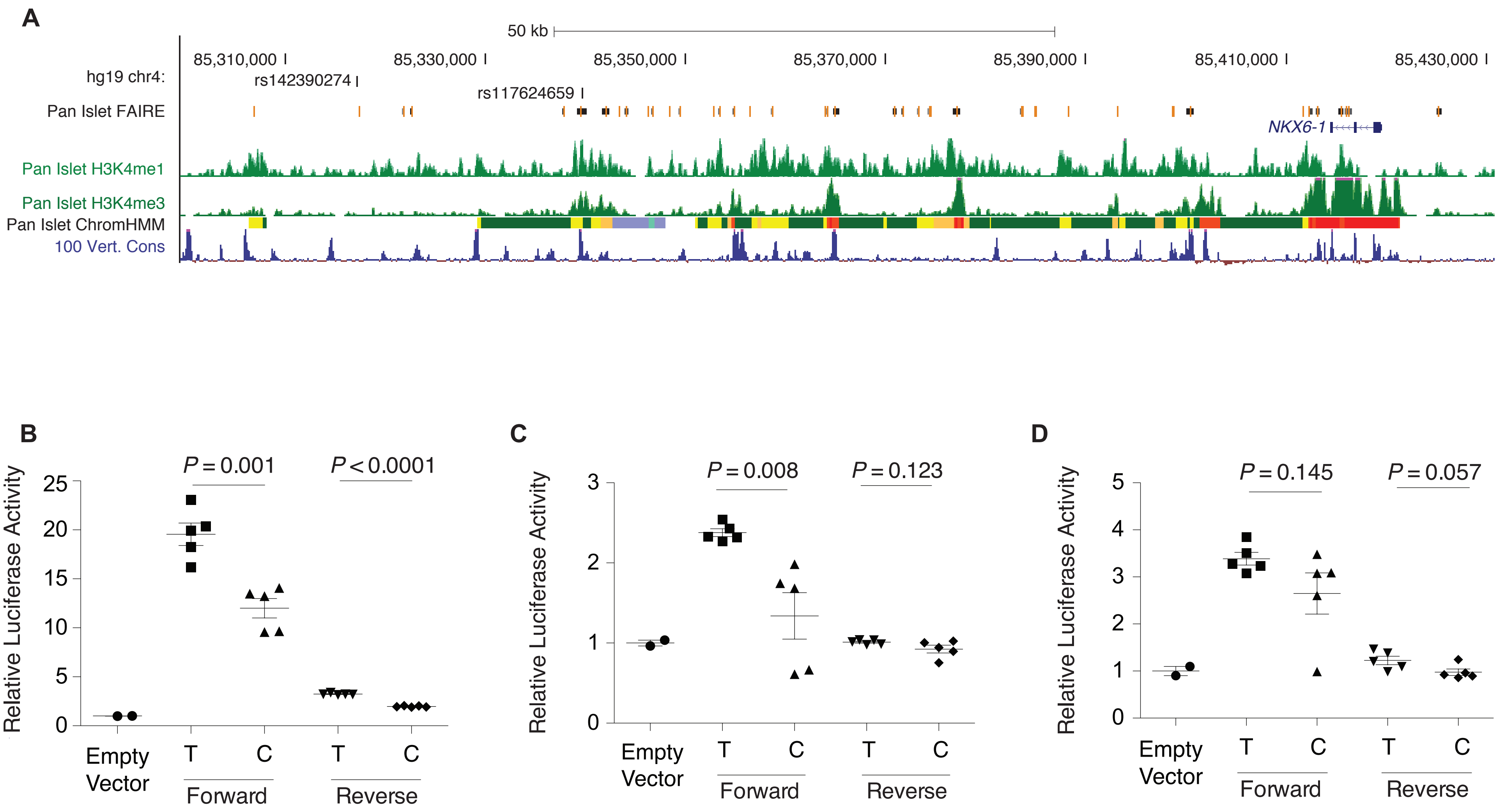

### Supplementary Figure 12

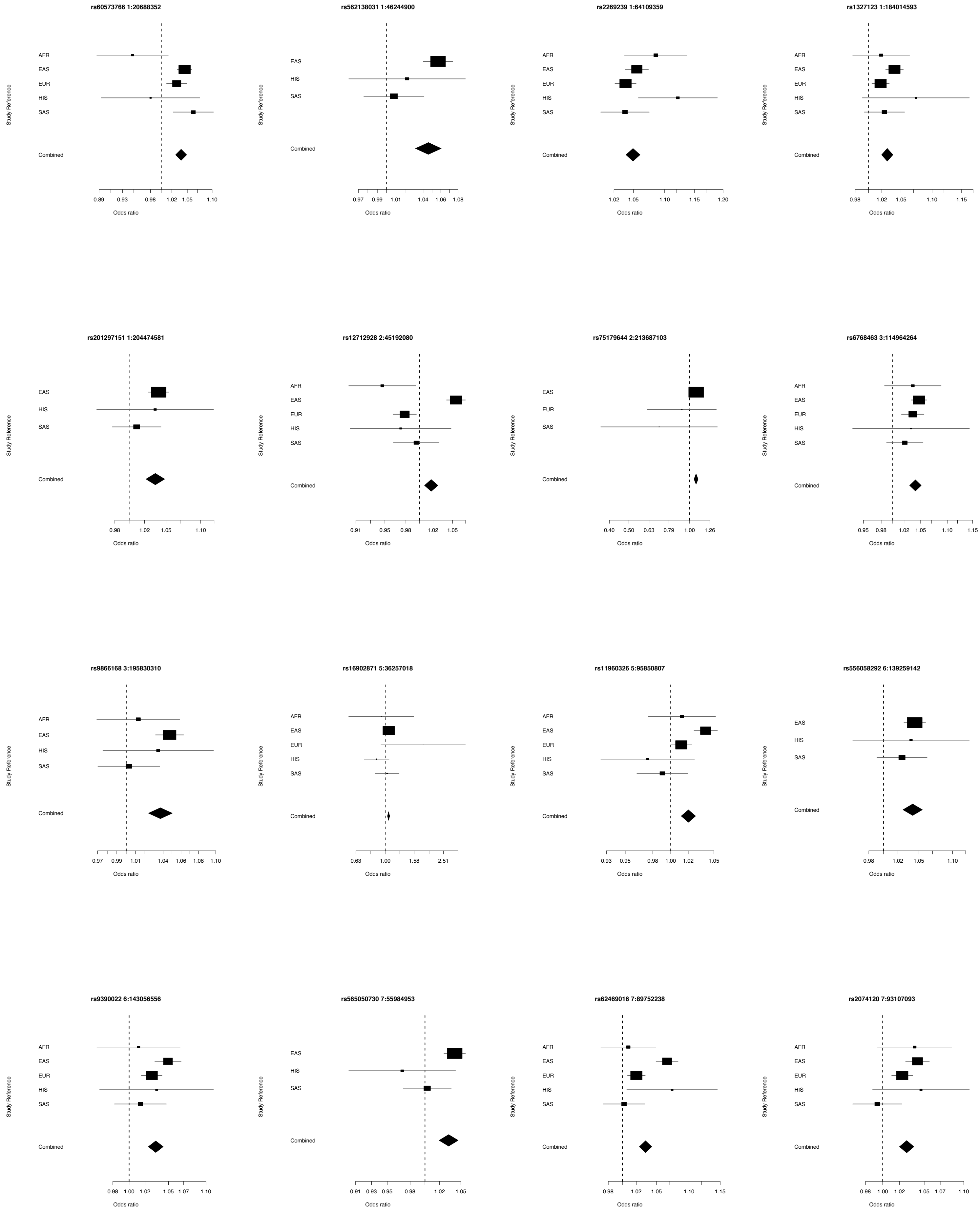

### Supplementary Figure 12, continued

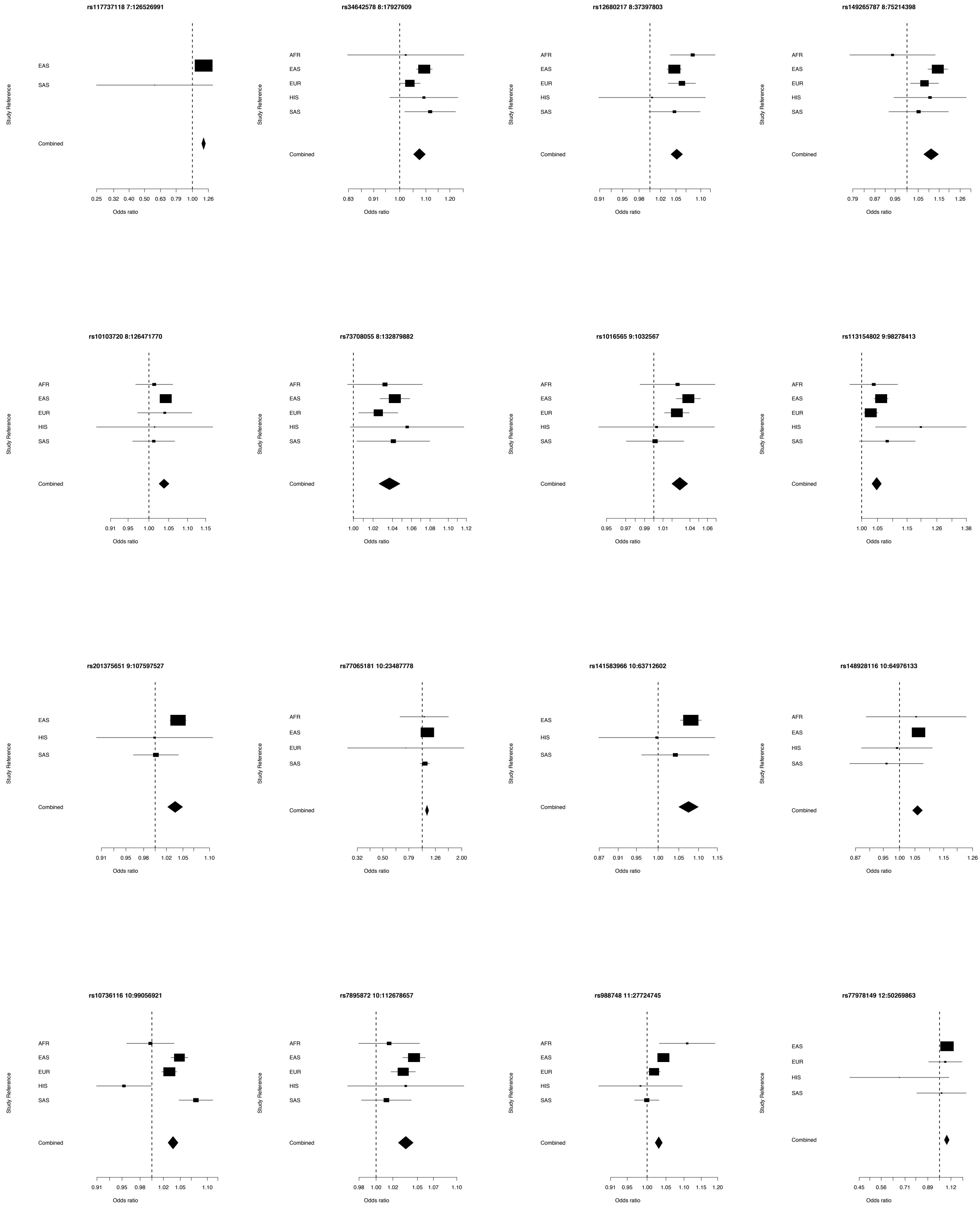

### Supplementary Figure 12, continued

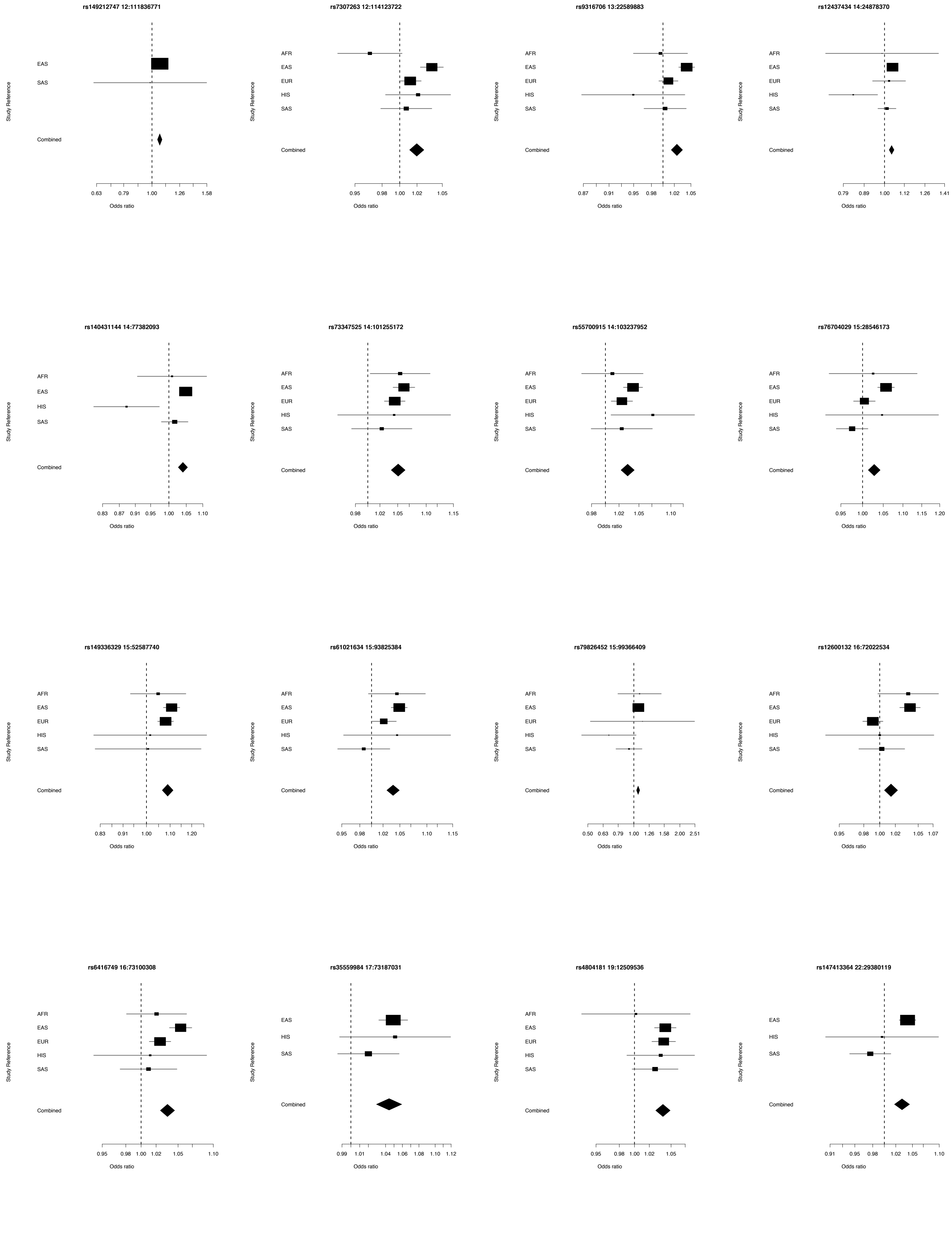

### Supplementary Figure 12, continued

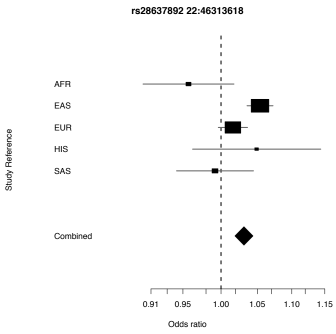

Supplementary Figure 13

**rs12712928**

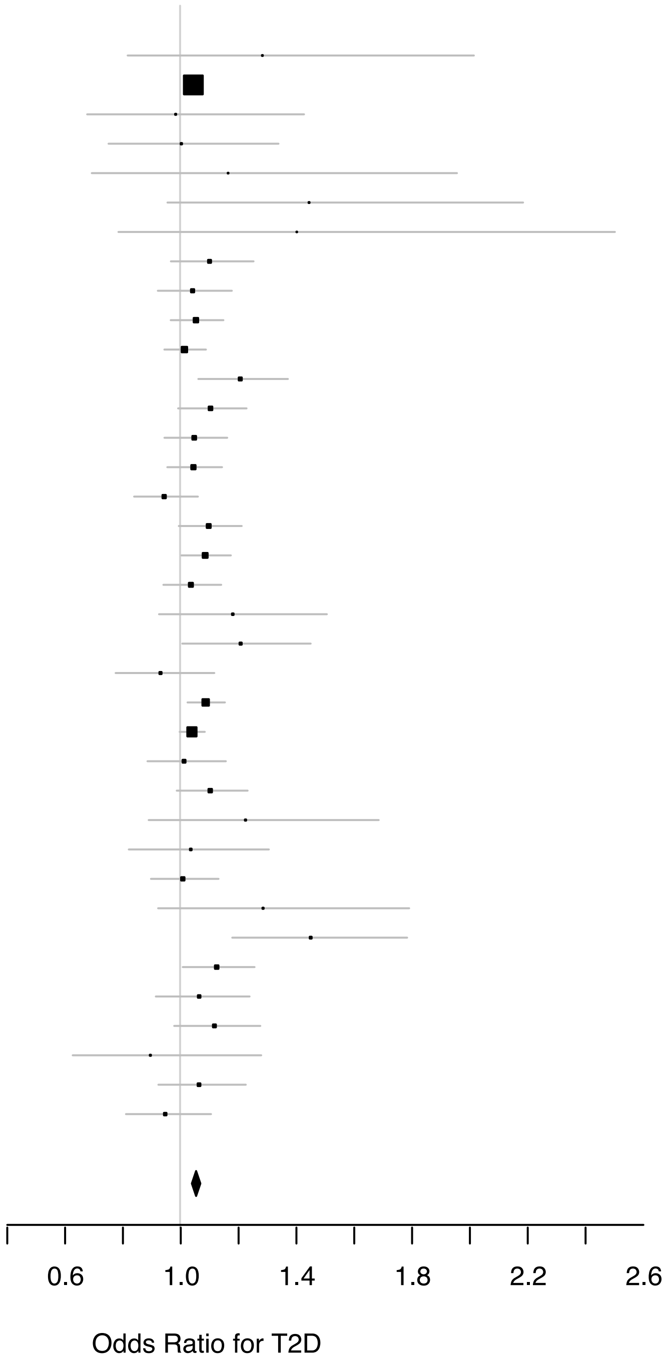


| STUDY | OR (95% CI) | P |
|---|---|---|
| AASC | 1.28 (0.82–2.02) | 2.78E−01 |
| BBJ | 1.05 (1.02–1.07) | 1.51E−05 |
| BES−1 | 0.98 (0.68–1.43) | 9.35E−01 |
| BES−2 | 1.00 (0.75–1.34) | 9.77E−01 |
| CAGE−1 | 1.17 (0.69–1.96) | 5.74E−01 |
| CAGE−3 | 1.45 (0.96–2.19) | 8.28E−02 |
| CAGE−Amagasaki | 1.40 (0.79–2.50) | 2.51E−01 |
| CAGE−GWAS | 1.10 (0.97–1.25) | 1.41E−01 |
| CHNS | 1.04 (0.92–1.18) | 4.99E−01 |
| CKB−12 | 1.05 (0.97–1.15) | 2.25E−01 |
| CKB−16 | 1.01 (0.95–1.09) | 6.83E−01 |
| CKB−26 | 1.21 (1.06–1.37) | 3.73E−03 |
| CKB−36 | 1.11 (0.99–1.23) | 6.80E−02 |
| CKB−46 | 1.05 (0.95–1.16) | 3.61E−01 |
| CKB−52 | 1.05 (0.96–1.15) | 3.32E−01 |
| CKB−58 | 0.94 (0.84–1.06) | 3.44E−01 |
| CKB−68 | 1.10 (1.00–1.21) | 6.12E−02 |
| CKB−78 | 1.09 (1.00–1.18) | 3.91E−02 |
| CKB−88 | 1.04 (0.94–1.14) | 4.53E−01 |
| CLHNS | 1.18 (0.93–1.51) | 1.76E−01 |
| DC−SP2−1 | 1.21 (1.01–1.45) | 4.05E−02 |
| DC−SP2−2 | 0.93 (0.78–1.12) | 4.49E−01 |
| HEXA | 1.09 (1.03–1.16) | 5.22E−03 |
| HEXA−CAVAS | 1.04 (1.00–1.09) | 5.97E−02 |
| HKDR | 1.01 (0.89–1.16) | 8.41E−01 |
| KARE | 1.10 (0.99–1.23) | 7.98E−02 |
| MESA | 1.23 (0.89–1.69) | 2.12E−01 |
| NAGAHAMA | 1.04 (0.82–1.31) | 7.60E−01 |
| SBCS−SWHS | 1.01 (0.90–1.13) | 8.74E−01 |
| SCES−1 | 1.29 (0.92–1.79) | 1.35E−01 |
| SCES−2 | 1.45 (1.18–1.79) | 3.94E−04 |
| SCHS | 1.13 (1.01–1.26) | 3.33E−02 |
| SiMES | 1.07 (0.92–1.24) | 4.08E−01 |
| SNUH | 1.12 (0.98–1.28) | 9.69E−02 |
| SSH | 0.90 (0.63–1.28) | 5.50E−01 |
| TaiChi−G | 1.07 (0.93–1.23) | 3.78E−01 |
| TWT2D | 0.95 (0.81–1.11) | 5.03E−01 |
| **Combined** | **1.06 (1.04–1.07)** | **3.16E−14** |
